# Site effects depth denoising and signal enhancement using dual-projection based ICA model

**DOI:** 10.1101/2023.04.26.538366

**Authors:** Yuxing Hao, Huashuai Xu, Mingrui Xia, Chenwei Yan, Yunge Zhang, Dongyue Zhou, Tommi Kärkkäinen, Lisa D. Nickerson, Huanjie Li, Fengyu Cong

## Abstract

Combining magnetic resonance imaging (MRI) data from multi-site studies is a popular approach for constructing larger datasets to greatly enhance the reliability and reproducibility of neuroscience research. However, the scanner/site variability is a significant confound that complicates the interpretation of the results, so effective and complete removal of the scanner/site variability is necessary to realize the full advantages of pooling multi-site datasets. Independent component analysis (ICA) and general linear model (GLM) based denoising methods are the two primary methods used to denoise scanner/site-related effects. Unfortunately, there are challenges with both ICA-based and GLM-based denoising methods to remove site effects completely when the signals of interest and scanner/site-related noises are correlated, which may occur in neuroscience studies. In this study, we propose an effective and powerful denoising strategy that implements dual-projection (DP) theory based on ICA to remove the scanner/site-related effects more completely. This method can separate the signal effects correlated with noise variables from the identified noise effects for removal without losing signals of interest. Both simulations and vivo structural MRI datasets, including a dataset from Autism Brain Imaging Data Exchange II and a traveling subject dataset from the Strategic Research Program for Brain Sciences, were used to test the proposed GLM- and ICA-based denoising methods and our DP-based ICA denoising method. Results show that DP-based ICA

## 1 Introduction

It is now common practice to pool multi-site magnetic resonance imaging (MRI) datasets to study brain biomarkers of neuroscience, neuropsychiatry, and neurology to promote rigor and reproducibility of results (Button et al., 2013; Eickhoff et al., 2016; Van Horn & Toga, 2009). However, combing multiple datasets does introduce site-related noise, which confounds effects of interest, and complicates the interpretation of the final results (Casey et al., 1998; Focke et al., 2011; Friedman et al., 2008; Pohl et al., 2016; Takao et al., 2011; Venkatraman et al., 2015; Vollmar et al., 2010; Wegner et al., 2008; Zivadinov & Cox, 2008). Site-related noises arises from differences in scanners manufacturers, field strengths, hardware, software, pulse sequences, quality control, and data quality across sites (Jovicich et al., 2009). It has been shown that differences in acquisition parameters and software and hardware upgrades during data collection using the same scanner have non-negligible effects on almost all image derived phenotypes from structural images (such as cortical surface and gray matter volume), diffusion weighted images (such as diffusion tensor image (DTI) measures), and functional MRI (fMRI) data (Groves et al., 2011; Li et al., 2020). Hence, effective removal or deconfounding of site-related variability from the MRI data is a critical step to ensure the accuracy and reproducibility of findings generated from combined datasets.

Several approaches have been proposed for harmonization of multi-site MRI data, including methods based on the general linear model (GLM) (Fennema-Notestine et al., 2007; Glover et al., 2012; Venkatraman et al., 2015), and data-driven unsupervised learning methods such as independent component analysis (ICA) (Chen et al., 2014; Li et al., 2020) and recently proposed deep learning methods (C Monte-Rubio et al., 2022; Dinsdale et al., 2021; Tian et al., 2022). Two of the most popular methods to eliminate or minimize the site-related noise are based on GLM and ICA (Chen et al., 2014). GLM-based denoising method is easy to implement and often used in multi-site MRI data studies to minimize the site-related noise, in this case, utilizes site/study variables as covariates of no interest in group-level GLM analysis to control for site-related effects. Fortin et al. (Fortin et al., 2017) have adapted a GLM-based technique called ComBat (Johnson et al., 2007), an empirical Bayesian method for data harmonization that is popular in the field of genetics, to remove unwanted variation induced by sites while preserving the signal-related variation in neuroimaging studies. ComBat has been applied to harmonize DTI measures (Fortin et al., 2017), cortical thicknesses and functional connectivity measures (Yu et al., 2018), magnetic resonance spectroscopy measures (T. K. Bell et al., 2022), and positron emission tomography (PET) outcomes (Orlhac et al., 2018) showing good performance for removing site effects. ICA is an unsupervised data-driven statistical method that factorizes or decomposes the image data into a set of statistically independent non-Gaussian components reflecting different signal sources that generate the measured imaging data, including those related to signals of interest and uninteresting or confounding noise and artifacts (Chen et al. 2014) that can be removed from the data to generated a denoised clean dataset for further analysis. In practice, ICA is applied to a single modality to identify noise-related components that are then removed by regressing them from the data. In this case, the ICA has been used to do a data driven estimation of the site/scanner-related covariates of no interest that are regressed out of the data, rather than creating covariates to model the site-scanner effects based on strong assumptions (e.g., regressors are used that assume a constant effect for each site/scanner, which ignores within-site/data to day variations in these effects). ICA is typically applied to denoise individual MRI modalities, however, our previous work (Li et al., 2020) proposed a denoising method for multi-modal imaging measures that implemented linked ICA (LICA) (Groves et al., 2011) as a novel approach to denoising scanner effects from multi-study data. LICA simultaneously decomposes the multi-modal imaging data (for example, structural plus diffusion MRI derived measures) into a set of multi-modal components and a set of subject loadings quantifying the strength of each multi-modal component in each individual, with components reflecting true signals of interest as multi-modal covariance patterns, as well as artifacts and variability related to uninteresting effects like scanner and site differences. We found that several of the resulting LICA components from an analysis of multi-study data with scanner effects were associated with scanner variations and that these patterns could be effectively regressed from the data to obtain denoised data relatively free from scanner effects. We showed that multi-modal ICA-based denoising was more effective at removing scanner-related effects compared with the conventional GLM and single-modality ICA denoising methods. The reason for its superior performance is that even though all three approaches involve regression to remove scanner effects from the data, the data-driven estimates of the scanner effects from LICA of multi-modal MRI data provided more accurate model of scanner effects than assuming a constant effect or estimating effects based on single modality ICA to use as nuisance covariates for denoising.

In the present study, we aim to address another limitation of current methods for denoising scanner/site effects, namely existing methods for denoising site/scanner effects ignore the possibility of correlations between these effects and the effects of interest. For the conventional GLM approach, site-related variables are included as covariates of no interest or may be regressed out of data prior to higher level statistical modeling, which may lead to removal of interesting signals that are correlated with scanner/site variables and to weaker specificity of denoising. While ComBat tries to preserve the signal-related variation when denoising scanner/site effects, similar to the conventional GLM approach, it also assumes a constant effect for all datasets collected from the same site or the same scanner state, thus also ignoring the day-to-day variations in scanner performance. While ICA and LICA can identify scanner/site effects that capture day to day variations in scanner performance (Li et al., 2020), both approaches are vulnerable to identifying components that reflect a mixture of signal and noise effects, rather than separating the effects into two separate components. In our previous work, to retain signals of interest, only components that were associated with scanner effects and not signals of interest (e.g., had subject loadings that correlated only with scanner variables and not variables of interest) were removed from the data while mixed components were retained as a conservative approach to denoising (Chen et al., 2014; Li et al., 2020). One possibility to address this limitation for ICA-based techniques are to run the ICA with several different model orders to try to identify a decomposition with stable pure noise components that are not mixed with signals of interest. In practice, it is challenging to do this as different mixtures may arise at different model orders such that it may not be possible to have full separation at any model order.

To solve this problem for ICA-based methods, we propose a new ICA with dual-projection (ICA-DP) technique for denoising scanner/site effects. In this study, we focus on single modality ICA, with extension to multi-modal ICA to be done in future work. For ICA-DP, mixed components from single-modality ICA are separated into a part related to signal only and a part related only to noise by applying a projection procedure. The noise effects extracted from the mixed components via the projection step are combined with the other ICA components that reflected only site/scanner variance are then removed from the data using a second projection procedure. Our new method is tested using simulated MRI data and in vivo multi-site datasets to assess the performance of ICA-DP as compared with conventional ICA and ComBat denoising methods.

## 2 Methods

### 2.1 Dual-projection denoising improvement

#### ICA-DP denoising

The ICA-DP denoising procedure is summarized in Fig. 1. ICA-DP is inspired by the dual-regression approach for projecting a participants fMRI data onto a set of spatial maps derived from ICA of multi-subject fMRI data to identify subject-level spatial maps corresponding to each group level component (Beckmann et al., 2009; Filippini et al., 2009; Nickerson et al., 2017). In the case of ICA-DP, “group” ICA maps are obtained from applying ICA to a subject-series of image derivatives such as spatial maps of fractional anisotropy or mean diffusivity, surface maps of cortical thickness or pial surface area, concatenated together into one 4D (N subjects x 3D volume/surface). This decomposition provides a set of spatial maps and a set of subject loadings, one for each component, that are then used for the dual projection procedure as follows.

**Fig. 1.**
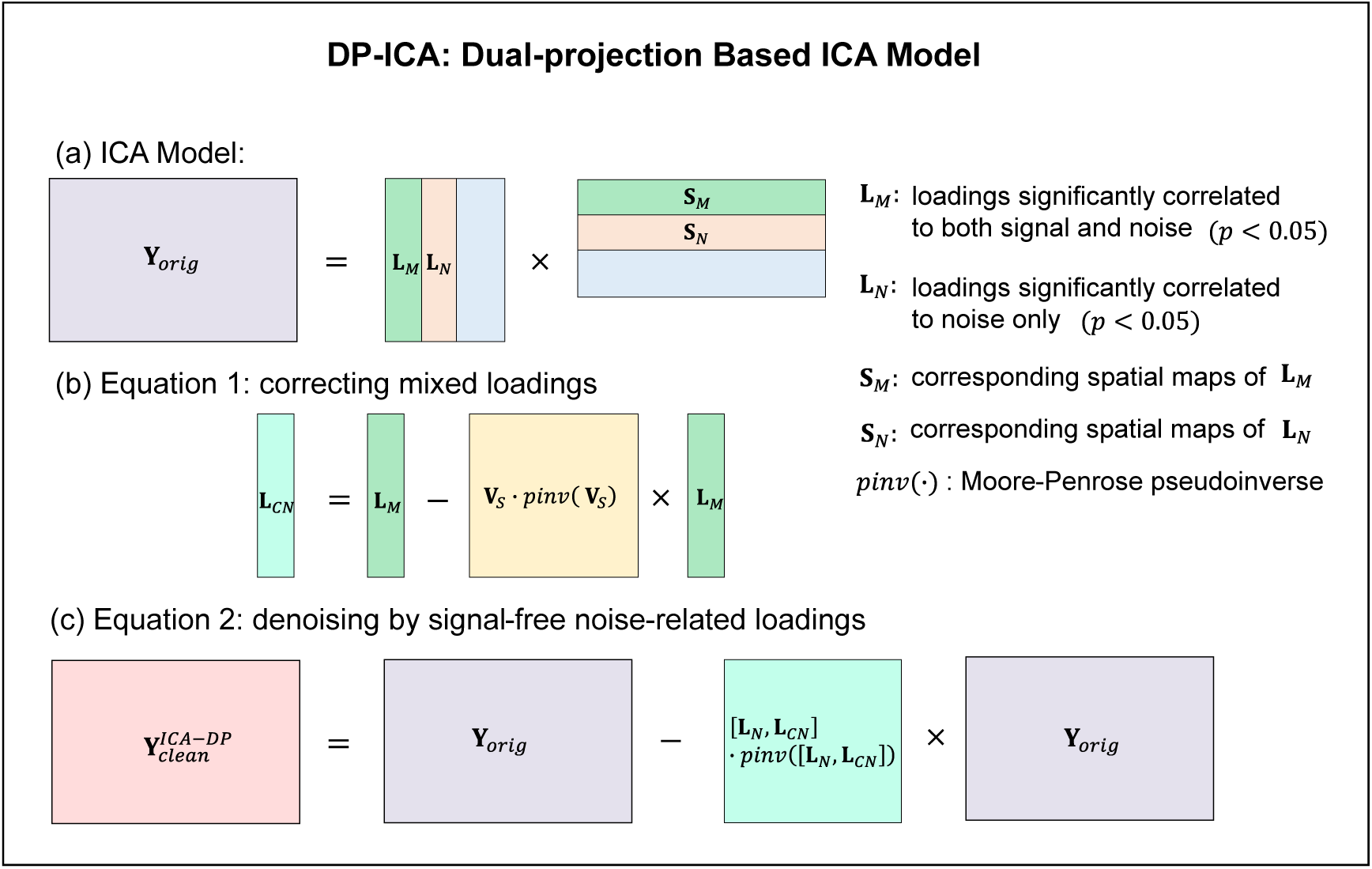
The procedures of ICA-DP denoising method. (a) Identifying the loadings extracted by ICA that related to noise variables (including mixed ones that both significantly correlated to both noise and signal, and the ones only significantly correlated to noise). (b) Correcting the mixed loadings to only noise-related ones (**L**_*CN*_) by projecting out signal-related information. (c) Obtaining cleaned data by removing the integral noise-related components (**L**_*N*_, **L**_*CN*_) out.

First, the subject series is decomposed by ICA, and the resulting subject loadings **L**, of each component that reflect the strength of the corresponding signal represented in the IC map (could be interesting signal, scanner/site/noise signals, or a mixture) are labelled as loadings for pure noise components **L**_*N*_, pure signal components **L**_*S*_, or mixed components **L**_*M*_ by calculating the correlations between the loadings and all the signal and noise variables (i.e. components exhibiting significant associations *p*<0.05 are identified as signal-and/or noise-related ones). The first projection procedure is used to separate the signal effects out from **L**_*M*_ as below:

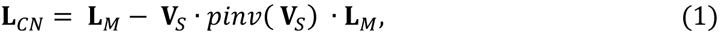

where **V**_*S*_ denotes the variables of interest (signals), **L**_*CN*_ denotes the corrected noise contributions to the mixed components, and *pinv*(·) denotes Moore-Penrose inverse (pseudoinverse) of a non-square matrix. Thus, the signal information is projected out from **L**_*M*_ and we can identify the site effects for the mixed components.

Then [**L**_*N*_, **L**_*CN*_] represent as the total site-related effects present in the data, which are then cleaned from the subject-series via a second projection procedure:

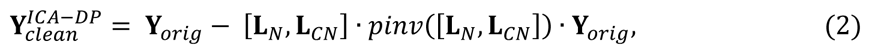

where **Y**_*orig*_ denotes the subject series of spatial maps and **Y**_*clean*_^*ICA-DP*^ denotes the denoised MRI data free from site/scanner effects that can be used for further analysis.

#### Previous denoising methods

##### ICA-SP denoising

When removing site effects of multi-site structural MRI data, the original ICA-based denoising methods (Chen et al., 2014; Li et al., 2020) only eliminate the pure noise components (only related with site effects) to avoid discarding useful information such as diagnoses or symptom measures. For comparison, we designate this ICA-based denoising method as ICA-SP (single projection) as it only uses one step of projecting:

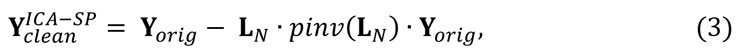

where **L**_*N*_ is the loadings of pure noise components and **Y**_*clean*_^*ICA-SP*^ is the denoised data derived from ICA-SP denoising methods.

Though it may preserve signal-related information well, it is too soft to remove the site effects as it does nothing with the mixed components and is more than likely to find site effects in its denoised data.

##### GLM-based denoising

The GLM model utilized for denoising MRI data is as follows (Maikusa et al., 2021; Venkatraman et al., 2015; Yamashita et al., 2019):

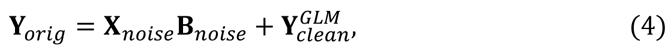

where **X**_*noise*_ is a design matrix for the site effects and **X**_*noise*_ is the regression coefficients corresponding to **X**_*noise*_(i.e., the spatial maps of site effects estimated by GLM model). In the original GLM model, **Y**_*clean*_^*ICA-DP*^ in equation (4) represents the model error. Here **Y**_*clean*_^*ICA-DP*^ is the cleaned data as the parameters in the design matrix represent site effects.

For denoising, the site variable in design matrix **X**_*noise*_ is coded as categorical variables, similar to the coding used in ANOVA factor effects models. For example, if the data **Y**_*orig*_ is from 3 different sites, then the site variable has 3 levels. For each coding variable, rows of the design corresponding to site *i* (*i* = 1,2,3) are set to 1 and others to 0. It is worth noting that GLM-based denoising is too hard when the signal and noise variables are significantly correlated, which will distort the signal-related information.

##### ComBat

Fortin et al. (Fortin et al., 2017) reformulated the ComBat model that Johnson et al. used in the context of gene expression analysis (Johnson et al., 2007) to DTI images:

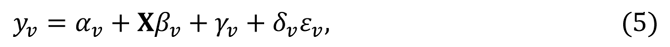

and reconstructed it as follows:

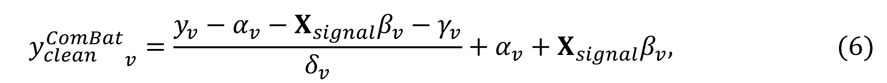

where **L**_*N*_ is the overall measure for voxel *v*, **X**_*noise*_ is a design matrix for the signal covariates like gender or age, and *²*_*v*_ is the voxel-specific vector of regression coefficients corresponding to **X**_*signal*_. The terms *γ*_*v*_ and *δ*_*v*_ represent the additive and multiplicative site effects of voxel *v*, respectively.

[**X**_*signal*_, **X**_*noise*_] was utilized (Fortin et al., 2017) to replace **X**_*signal²*_*v* + *Y*_*v*_ (MATLAB codes: https://github.com/Jfortin1/ComBatHarmonization) to estimate the parameters in (6) and add the signal-related part **X**_*signal*_,**β**_*v*_ back after removing the site effects, thus can alleviate the signal distortion in the GLM-based denoising procedure.

Different from the data-driven denoising methods based on ICA, the denoising schemes based on GLM and ComBat are model-based that the noise is constructed by users rather than extracted from the data.

### 2.2 Study Data

#### Simulated Data

The simulated structural MRI data including 100 subjects were generated in this study. For each subject, the data was generated by computing 10 spatially maps and one set of ground truth subject loadings. Each component map was multiplied by the corresponding subject loading, and then they were added together to obtain the simulated MRI data for each subject. The spatial maps were gotten by combining different areas of the standard brain template as shown in Fig. 2 (https://github.com/ThomasYeoLab/CBIG/tree/master/stable_projects/brain_parcellation/Schaefer2018_LocalGlobal/Parcellations/). To make our simulated data much close to real MRI data, two kinds of spatial maps were simulated, one is all the spatial maps of 10 components were spatially independent and the other is two components’ spatial maps were overlapped (Fig. 2). For each condition, 100 subjects were generated, and the subject-specific data shared the same spatial maps, and the difference was the weights in its loadings corresponding to the spatial maps. Three different types of relationships between subject loadings and signal/noise variables were simulated in this study: (1) signal variable was not significantly correlated to noise variable; subject loadings were linearly correlated to signal and (or) noise variables (Table 1). Among the 10 components, the first four components were significantly related to signal and (or) noise variables, and the other components were not related to the variables we are interested in. Components 1 and 2 are mixed components, which are related with both signal and noise variables. The difference is that component 1 is much more related to signal, and component 2 is much more correlated to noise. Component 3 is pure noise components, which only significantly correlated with noise variable. Component 4 is pure signal component that only significantly correlated with signal variable; (2) Signal variable is significantly correlated to noise variable, subject loadings are linearly correlated to signal and (or) noise variables (Table 2). Since signal variable was significantly correlated to noise variable, there were no pure signal or noise components under this condition, so we selected the first 2 components as mixed components, component 1 is much more related to signal, and component 2 is much more correlated to noise. Three different correlation levels (from low to high) between signal and noise variables were simulated in this study to show the denoising power of ICA-DP. (3) Signal variable and noise variable were not correlated, but the subject loadings were non-linearly correlated to noise variable. For this case, we chose the first component as the pure noise component, whose loading is correlated with the element-wise square of the noise variable, the correlation level is 0.8000. The other components were non-interested components.

**Fig. 2.**
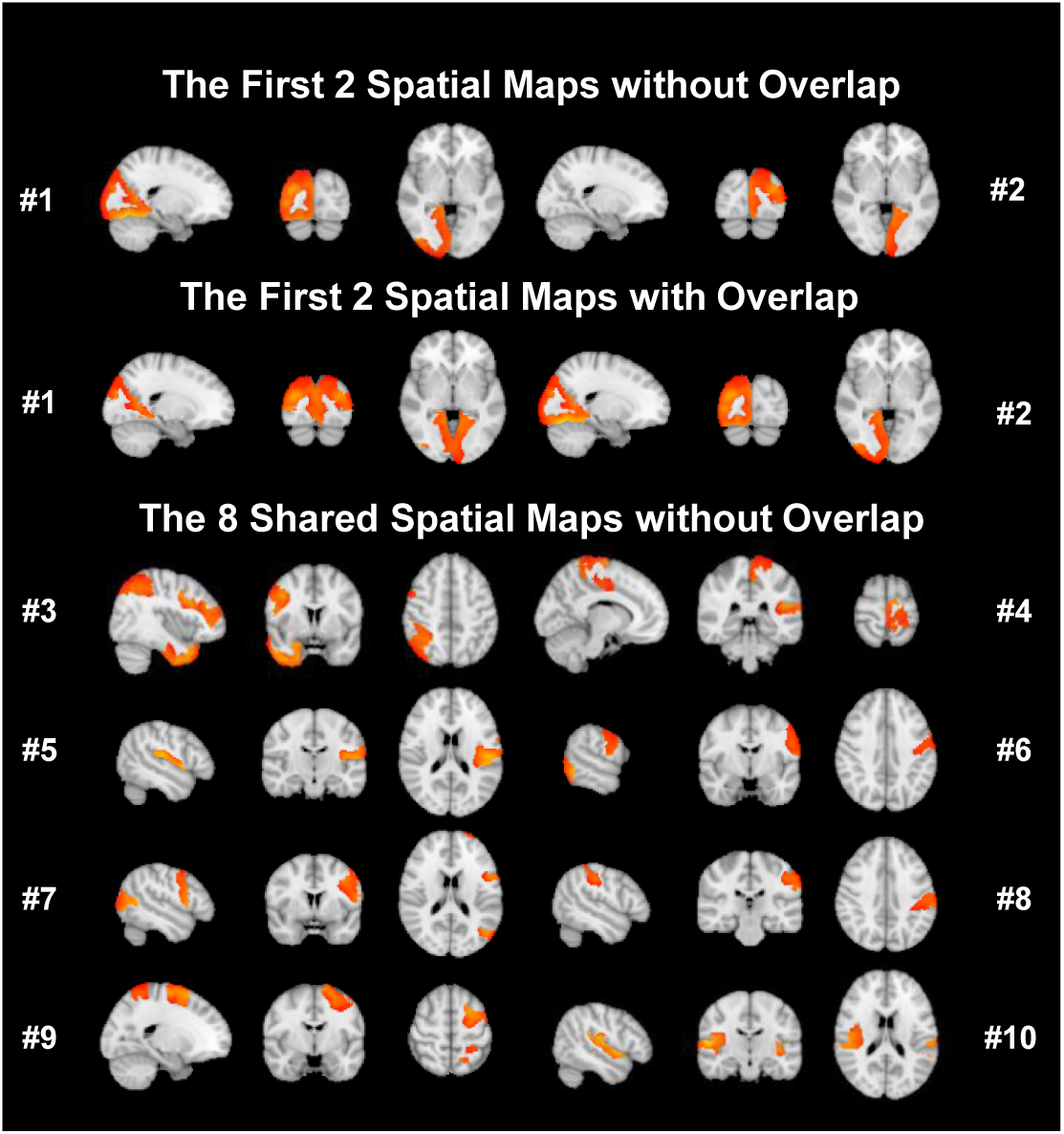
Ten independent brain spatial maps used to simulate MRI data. Situation1: there are no overlap among all the spatial maps; Situation 2: The first two components were spatially overlapped, and the other 8 components shares the same maps with situation 1.

**Table 1.**
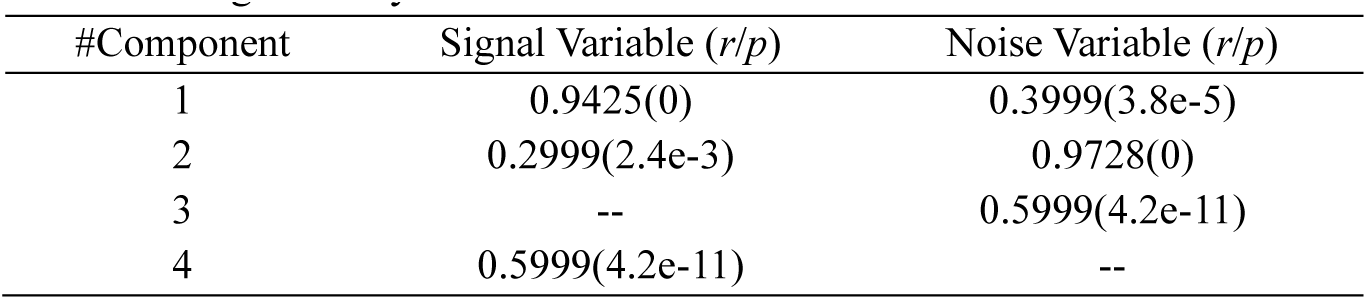
Component loadings are linearly correlated with signal and noise variables, while signal variable is not significantly correlated to noise variable. Components 1 (more related to signal) and 2 (more related to noise) are mixed components. Component 3 is only related with noise variable and component 4 is only related with signal variable. The relationship of loadings and variables is expressed by *r*-value and *p*-value. -- denotes not significantly correlated.

**Table 2.** Component loadings are linearly correlated with signal and noise variables, while signal variable is significantly correlated to noise variable. Components 1 (more related to signal) and 2 (more related to noise) are mixed components. The relationship of loadings and variables is expressed by *r*-value and *p*-value. Three different correlation levels between signal variable and noise variable are simulated in this study with *r-*value of 0.2999 (*p*=0.0024), 0.4999 (*p*=1.2e-7), and 0.6999 (*p*=5.6e-16), corresponding to low, medium and high correlation levels.

| Correlation between signal and noise | #Component | Signal | Noise |
| --- | --- | --- | --- |
| 0.2999 (2.4e-3) | 1 | 0.7946(0) | 0.2412(1.6e-2) |
|  | 2 | 0.2590(9.3e-3) | 0.7959(0) |
| 0.4999 (1.2e-7) | 1 | 0.7962(0) | 0.4279(8.9e-6) |
|  | 2 | 0.3447(4.5e-4) | 0.7859(0) |
| 0.6999 (5.6e-16) | 1 | 0.7957(0) | 0.5932(7.8e-11) |
|  | 2 | 0.4993(1.2e-7) | 0.7761(0) |

#### Multi-site MRI data from ABIDE II

High spatial resolution structural MRI data of 606 subjects (including Autism Spectrum Disorder (ASD) patients: 225, Healthy Controls (HC): 381) were obtained from Autism Brain Imaging Data Exchange II dataset (ABIDE II) (http://fcon_1000.projects.nitrc.org/indi/abide/abide_II.html). The data were collected from 13 different sites, all the data were acquired from 3T scanner with different manufacturers (Siemens, Philips, and GE) (Di Martino et al., 2017). The acquisition parameters: scanner and imaging-related details, including repetition time (TR), echo time (TE), flip angle (FA), voxel size in Table 3. And the demographic information: ASD/HC, gender and age are summarized in Table 4. Subjects with ASD could be divided into two categories: ASD only and ASD with comorbidity (Attention-Deficit/Hyperactivity Disorder, anxiety or orthers) (Di Martino et al., 2017).

**Table 3.** Scanning parameters and demographic information of the multi-site ABIDE II data. The data were collected from 13 different sites: Erasmus University Medical Center (EMC), ETH Zürich (ETH,), Georgetown University (GU), Indiana University (IU), Kennedy Krieger Institute (KKI), Katholieke Universiteit Leuven (KUL), Oregon Health and Science University (OHSU), Olin Neuropsychiatry Research Center (ONRC), Stanford University (SU), University of California Davis (UCD), University of California Los Angeles (UCLA), University of Miami (UM), University of Utah School of Medicine (USM).

| Sites | Scanners | TR/TE (ms) | FA (degree) | Voxel Size |
| --- | --- | --- | --- | --- |
| EMC | GE MR750 | 1664/4.24 | 16 | 0.9×0.9×0.9 |
| ETH | PhilipsAchieva | 3000/3.9 | 8 | 0.9×0.9×0.9 |
| GU | Siemens TriTim | 2530/3.5 | 7 | 1×1×1 |
| IU | Siemens TriTim | 2400/2.3 | 8 | 0.7×0.7×0.7 |
| KKI | PhilipsAchieva | 3500/3.7 | 8 | 1×1×1 |
| KUL | PhilipsAchieva | 2000/4.6 | 8 | 1×1×1.2 |
| OHSU | Siemens TriTim | 2300/3.58 | 10 | 1×1×1.1 |
| ONRC | Siemens Skyra | 2200/2.88 | 13 | 0.8×0.8×0.8 |
| SU | GE SIGNA | 5.9/1.8 | 11 | 1×1×1 |
| UCD | Siemens TriTim | 2000/3.16 | 8 | 1×1×1 |
| UCLA | Siemens TriTim | 2300/2.86 | 9 | 1×1×1.2 |
| UM | GE Healthcare | -/- | 12 | 1×1×1 |
| USM | Siemens TriTim | 2300/2.91 | 9 | 1×1×1.2 |

**Table 4.**
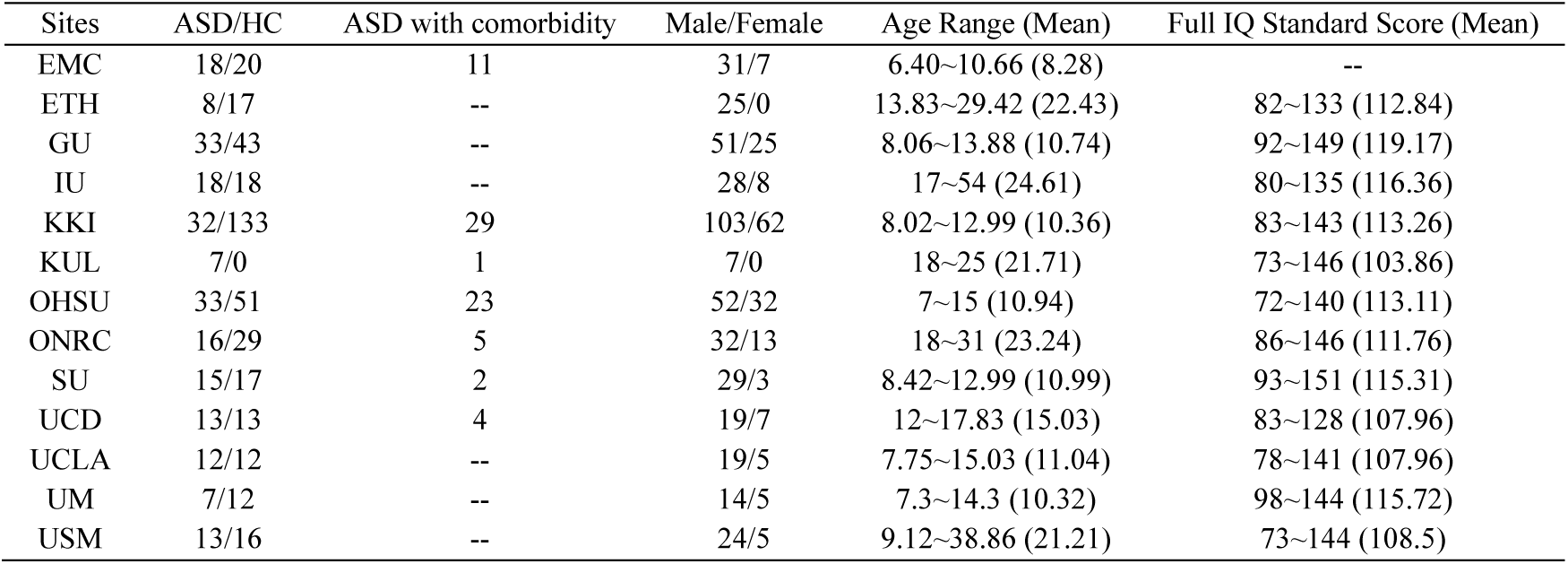
Demographic information of the multi-site ABIDE II data collected from 13 different sites.

In this study, the site differences are defined as noise variables, while group difference (ASD/HC), age and gender are regarded as signal variables. The correlation coefficients among these variables are summarized in Table 5. Since site difference are categorical variables and calculating the correlation coefficients between categorical variable and numeric variable directly is not achievable, we used ANOVA to calculate the significant levels of signal variables and site noise variable. For ICA analysis, the components that only significantly correlated to site variables were regarded as pure noise components and the components significantly correlated to both site and signal variables were regarded as mixed components.

**Table 5.** The relationship of signal and noise variables for real MRI data.

| Correlation | Site | ASD vs HC | Age | Gender |
| --- | --- | --- | --- | --- |
| Site | 1.000(0.0000) | 4.8e-6 | 1.3e-160 | 0.0004 |
| ASD vs HC | 4.8e-6 | 1.000(0.0000) | - | 0.1959(1.2e-6) |
| Age | 1.3e-160 | - | 1.000 (0.0000) | - |
| Gender | 0.0004 | 0.1959(1.2e-6) | - | 1.000(0.0000) |

#### Traveling subjects

To further validate the site effects removing efficiency of ICA-DP, the high spatial resolution structural MRI data from a traveling-subject dataset including 9 healthy participants (all males, age: 27±2.6) scanned at 12 different sites from the DecNef Project Brain Data Repository (https://bicr-resource.atr.jp/srpbsts/) were used in this study. For T1-weighted MRI data of the 12 different sites, there were two phase-encoding directions (PA and AP), three MRI manufacturers (Siemens, GE, and Philips) with seven scanner types (TimTrio, Verio, Skyra, Spectra, MR750W, SignaHDxt, and Achieva) and four channels per coil (8, 12, 24, and 32) (Maikusa et al., 2021; Tanaka et al., 2021). Scanning parameters including repetition time (TR), echo time (TE), flip angle (FA) and voxel size, are summarized in Table 6. Three sites (i.e., ATT, UTO and YC2) were excluded for denoising analysis because they contain duplicate data.

**Table 6.** Scanning parameters of the traveling-subject dataset. The datasets include 9 healthy subjects undergoing T1-weighted MRI scans at 12 different sites, and all of them used 3T scanners and same acquisition parameters but with different manufacturers and hardware versions (Siemens, GE, and Philips).

| Sites | Scanners | TR/TE (ms) | FA (degree) | Voxel Size |
| --- | --- | --- | --- | --- |
| ATT | SiemensTimTrio | 2300/2.98 | 9 | 1×1×1 |
| ATV | Siemens Verio | 2300/2.98 | 9 | 1×1×1 |
| COI | Siemens Verio | 2300/2.98 | 9 | 1×1×1 |
| HKH | Siemens Spectra | 1900/2.38 | 10 | 0.8×0.75×0.75 |
| HUH | GE Signa HDxt | 6788/1.928 | 20 | 1×1×1 |
| KPM | Philips Achieva | 7.1/3.31 | 10 | 1×1×1 |
| KUS | Siemens Skyra | 2300/2.98 | 9 | 1×1×1 |
| KUT | SiemensTimTrio | 2000/3.4 | 8 | 0.9375×0.9375×1 |
| SWA | Siemens Verio | 2300/2.98 | 9 | 1×1×1 |
| UTO | GE MR750W | 7.7/3.1 | 11 | 1×1.0156×1.0156 |
| YC1 | Philips Achieva | 6.99/3.176 | 9 | 1×1×1 |
| YC2 | Philips Achieva | 7.01/3.155 | 9 | 1×1×1 |

### 2.3 Data analysis

For both ABIDE II dataset and travelling dataset, all the data were processed based on FMRIB Software Library, FSL (https://fsl.fmrib.ox.ac.uk/fsl/fslwiki/). Grey matter (GM) images were generated with high-spatial resolution structural MR images with FSL-VBM (https://fsl.fmrib.ox.ac.uk/fsl/fslwiki/FSLVBM). Firstly, non-brain tissue was removed, followed by GM-segmentation. GM images were non-linearly registered to MNI 152 standard space. The resulting images were concatenated and averaged to create a study-specific grey matter template. Then, all native GM images were non-linearly registered to the study-specific template. Modulated GM images were then smoothed using an isotropic Gaussian kernel (sigma = 3mm).

### 2.4 Denoising procedures and assessments

#### Site effects removing with ICA-SP/DP methods

Firstly, ICA was applied to the non-denoised data to identify pure noise components, pure signal components and mixed components. For simulated data, the Pearson correlation coefficient between subject loadings and signal (or noise) variable was used as a measure of the properties of components. Those components only related with the signal variable (*p* < 0.05) were identified as pure signal components; those components only related with the noise variable (*p* < 0.05) were identified as pure noise components; those components related with the signal and noise variable (*p* < 0.05) were identified as mixed components.

For the real MRI data from ABIDE II and the traveling-subject data, we used the Pearson correlation coefficient and ANOVA to identify signal, noise and mixed components (age, gender, and group difference(ASD/HC) were signal variables of interest). The subject loadings from one site were divided into one variable, then 13 levels-for ABIDE II and 9 levels-for the traveling-subject data ANOVA were used to calculate the significant levels of subject loadings and site difference. Finally, those components whose p-values of ANOVA were significant (using Bonferroni correction to adjust for multiple comparisons, adjusted *p* < 0.05) were identified as site-related components. The intersection of signal components and site-related components were mixed components.

For simulated data, the data were decomposed into 10 components based on the simulation. For the data from ABIDE II, they were decomposed into 50, 100 and 150 independent components, by calculating the correlation levels of subjects’ loadings and variables with the analysis of variance (ANOVA) for each component, we identified the numbers of pure noise components were 27, 56 and 96, respectively, and the numbers of mixed components were 23, 42 and 49, respectively. For the traveling-subject data, they were decomposed into 50 independent components, and 10 pure noise components and 16 mixed components were identified.

For ICA-SP method, only pure noise components were used to regress out the site effects. For ICADP method, all the noise-related components, including the mixed ones, were used for site effects removal. The noise effects extracted from the mixed components and pure noise components will be regarded as the integral site-related noise effects for removal by ICA-DP. All the ICA analysis was based on MATLAB and FSL MELODIC.

#### Site effects removing with GLM and ComBat methods

For GLM-based denoising method, the site difference is regarded as the covariates to be regressed out. For ComBat denoising method, firstly, ComBat normalizes the data by removing the effect of the overall mean and signal variables. Then using an empirical Bayes framework, ComBat gets an estimation for additive and multiplicative site effects. The final denoised data could be obtained by removing these noise effects and adding the signal-related information back. In our study, the performance of GLM was only evaluated with simulation data, the main reason is that ComBat is a GLM-derived model and more powerful than the original GLM model. Thus, for real MRI data, only the performance of ICA model and ComBat model were compared.

#### Evaluating the denoising results

For simulated MRI data, ICA was utilized to the non-denoised and denoised data to extract and identify the signal-and noise-related components to compare the denoising effects of all the methods.

For the real MRI data, a set of analyses were applied to show the performance of site effects elimination and biological variability preservation (including age effects and group difference (ASD/HC)) for all the denoising methods. For site effects removing evaluation, T-distributed stochastic neighbor embedding (t-SNE) was used to visualize the heterogeneity related with sites of non-denoised data and denoised data by projecting their dominant features into a 2D space. We could assess the efficiency of the denoising methods by checking whether there are site-clustered distributions or noticeable inter-site heterogeneities after site effects removing. Group F-test was also applied to both the non-denoised data and the denoised data derived from different methods to test the significant difference regions caused by the site difference. For age effects evaluation, the Pearson correlation coefficient between median GM and age of all the subjects from ABIDE II data was calculated to show the relationship between them. The median GM per subject was obtained by calculating the median of 100 regions of interest (ROI), which is the template used to generate the simulated MRI data. Group t-test was also applied to both the non-denoised data and the denoised data derived from different methods to test the significant difference regions caused by the age. For group difference (ASD/HC) evaluation, group t-test was applied to both the non-denoised data and the denoised data derived from different methods to test the significant difference regions caused by the group difference. FWE-corrected *p* < 0.05 using non-parametric permutation testing with threshold-free cluster enhancement (TFCE) (Smith & Nichols, 2009) in FSL’s Randomise (Winkler et al., 2014), with 5,000 permutations was used to find the significant regions.

## 3 Results

### 3.1 Simulation Denoising Results

Fig. 3 shows the signal-and noise-related components of simulated data extracted by ICA, before and after denoising, when the signal variable is not significantly correlated to the noise variable. The results are shown in Fig. 3(a) when the spatial maps of all 10 components are spatially independent. Fig. 3(b) shows the results when the first two components are spatially overlapped. When the signal variable is not correlated to noise variable, the results for spatially independent data and spatial dependent data are similar. All the denoising methods could remove pure noise component #3 and preserve pure signal component #4. However, the noise effects cannot be denoised from the mixed components #1 (more related to signal for the original data) and #2 (more related to noise for the original data) by ICA-SP method. The performance of ICA-DP, GLM and ComBat were comparable under this situation, the noise effects in the mixed components #1 and #2 were effectively removed and the signal effect was enhanced by increasing its correlation levels with the signal variable after denoising by ICA-DP, GLM and ComBat. The two mixed components were combined into one component that is only significantly related with signal variable. The noise-related regions were also removed after denoising with ICA-DP, GLM, and ComBat methods.

**Fig. 3.**
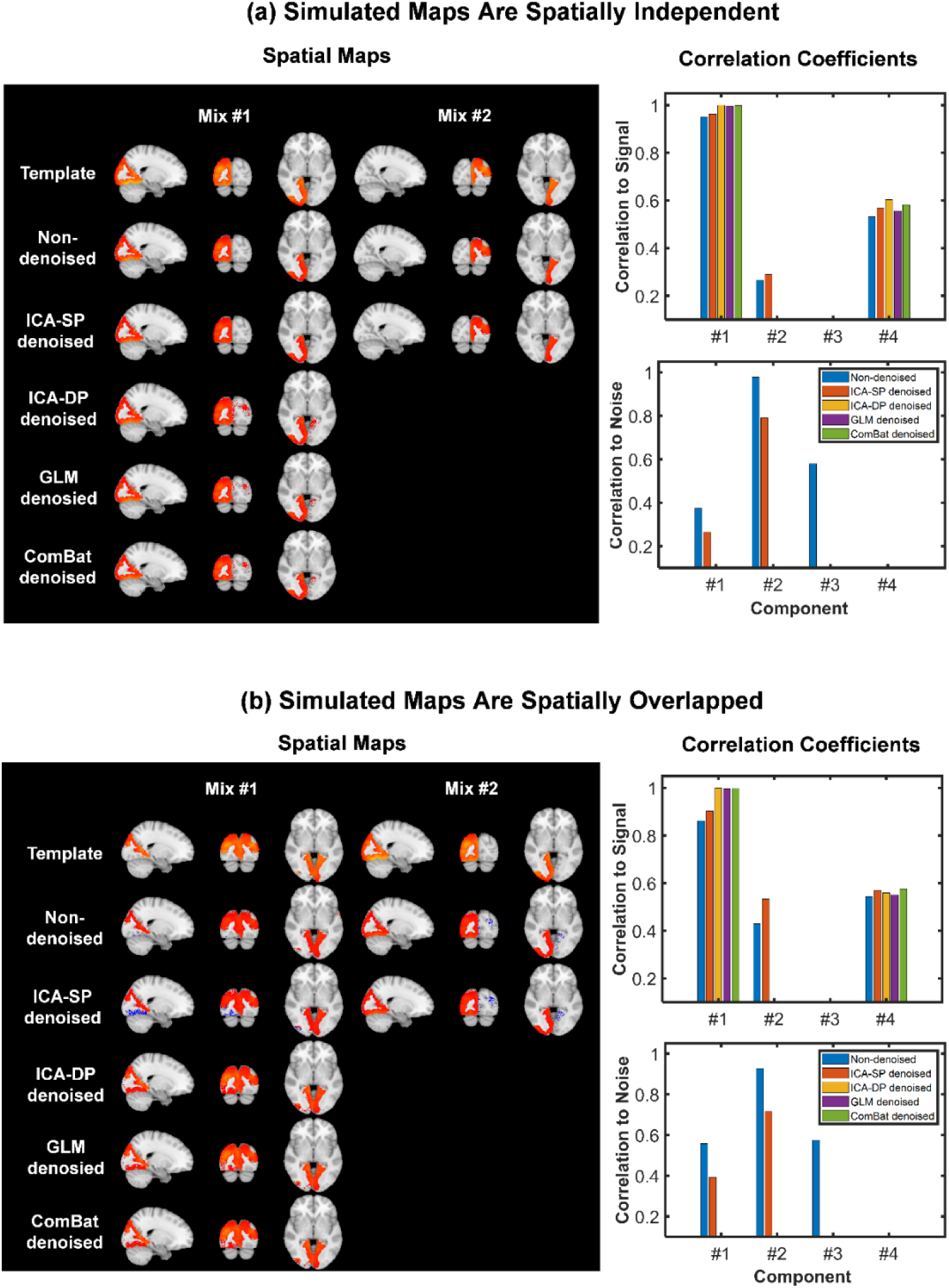
Denoising effects on the signal-and noise-related components when the signal variable is not significantly correlated to the noise variable. (a) Results when the spatial maps of all 10 components were spatially independent. (b) Results when the spatial maps of the first two components were spatially overlapped. For the non-denoised data, components #1 and #2 are two mixed components, named as Mix #1 (more related to signal variable) and Mix #2 (more related to noise variable), component #3 is pure noise component and component #4 is pure signal component. The pure noise component was removed by all the denoising methods. The two mixed components were combined into one component that is only significantly related with signal variable. The noise-related regions were also removed after denoising with ICA-DP, GLM and ComBat methods. ICA-SP cannot denoise the noise effects from the two mixed components.

Fig. 4 shows the denoising effects on the two mixed components when the signal variable is significantly correlated to the noise variable. Three different correlation levels between the signal and noise variables were simulated. Fig. 4(a) shows the results when the spatial maps of all 10 components were spatially independent. Fig. 4(b) shows the results when the spatial maps of the first two components were spatially overlapped. Among all the denoising methods, only ICA-DP could effectively weaken the noise while strengthening the signal effects when signal and noise variables are correlated for two types of simulated data.

**Fig. 4.**
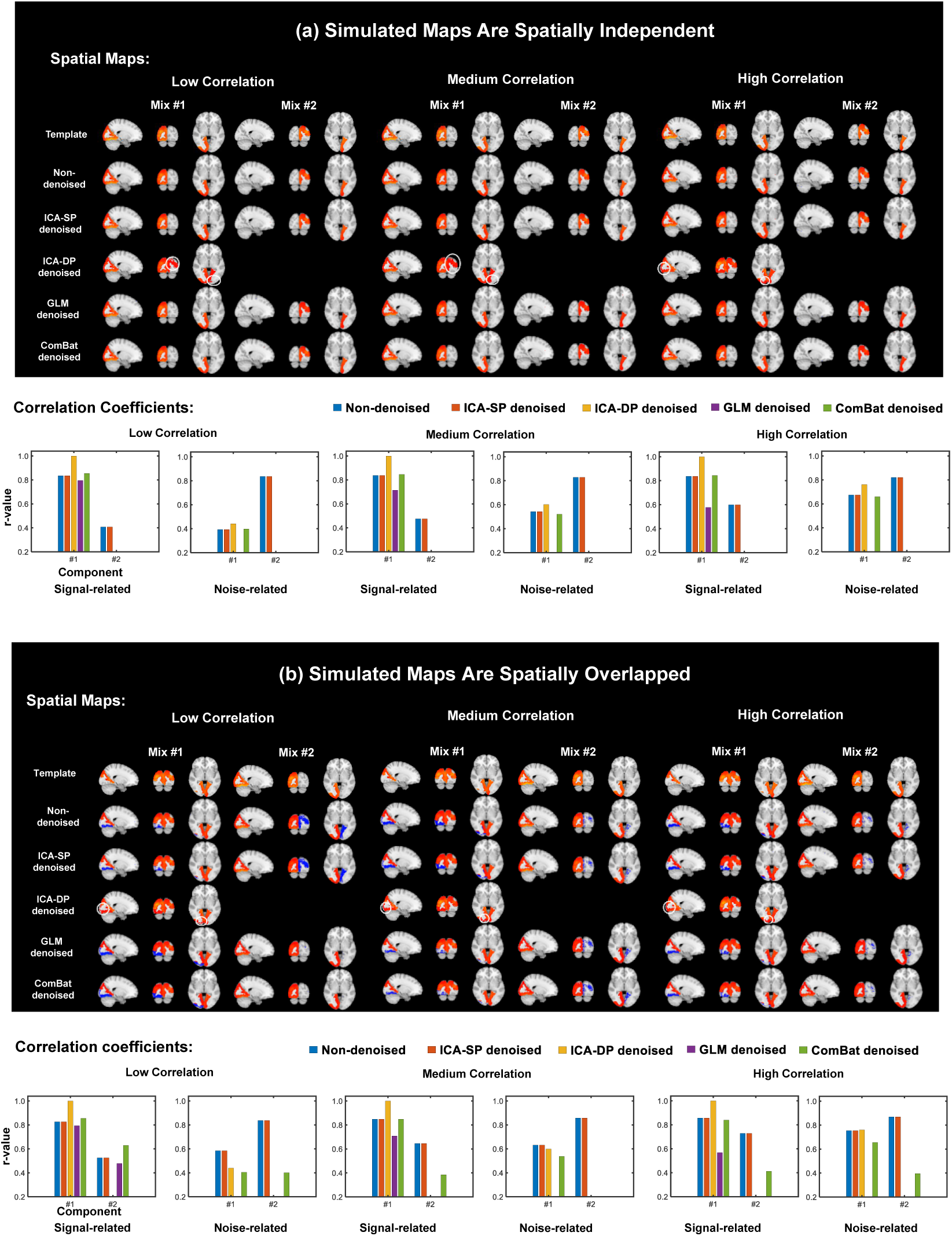
Denoising effects on the two mixed components when the signal variable is significantly correlated to the noise variable. Mix #1 is more related to signal variable and Mix #2 is more related to noise variable for the non-denoised data. Three different correlation levels between signal and noise variables were simulated in this study to test the denoising performance of all the denoising methods. (a) When the spatial maps of all 10 components were spatially independent. (b) Results when the spatial maps of the first two components were spatially overlapped. After denoising by ICA-DP, Mix #1 and #2 were merged into a single component that was more related to signal variable and the site-related effects were effectively decreased. Some spatial area that related with noise effects were also removed (highlighted with white circle), especially for lower correlation between signal and noise variables. Among all the denoising methods, only ICA-DP could effectively weaken the noise while strengthening the signal effects when signal and noise variables are correlated.

ICA-SP denoising could not remove the noise from the mixed components, as both the extracted spatial maps and correlation coefficients between subject loadings and noise variables of the mixed components were not changed after denoising by ICA-SP, compared to that of non-denoised data.

GLM denoising showed the most aggressive denoising performance while eliminating the noise-related information at the expense of destroying the signal-related information. After denoising by GLM, both components #1 and #2 were not correlated to noise variable. Besides, component #2 was not correlated to signal variable any longer and the correlation between component #1 and signal variable also became lower, which became more serious with the increasing of the correlation levels between signal and noise variables.

Combat denoising could not remove the noise effects when the mixed component is more related with signal effects (component #1) and showed aggressive denoising performance that also removes signal effects when the mixed component was more correlated with noise variable (component #2). When all the 10 components are spatially independent, after denoising by ComBat, both spatial maps and subject loadings of the mixed component #1 were not changed, while the mixed component #2 was not related with both signal and noise variables any longer. When the two mixed components are spatially overlapped, the noise effects could not be effectively removed by ComBat when the mixed component was more related to signal variable (component #1) and showed aggressive denoising performance that removed some signal effects when the mixed component was more related with noise effects (component #2). Both spatial maps and subject loadings of the mixed component #1 (more related to signal) did not change significantly, while the mixed component #2 (more related to noise) was less correlated to both signal and noise variables.

After denoising by ICA-DP, the original mixed components #1 and #2 were merged into a single component that was more related to signal variable and the site-related effects were effectively decreased. Some spatial area that related with noise effects were also removed (highlighted with white circle), especially for lower correlation between signal and noise variables. Fig. 4(b) shows that the removed spatial parts only cover the unique parts of component #2 and do not involve the overlapping parts. Though the merged component after denoising by ICA-DP was still mixed component, the correlation coefficient between its loading and signal variable was strengthened for all the signal to noise correlation levels, and the correlation levels to noise variable were contributed by the inherent relationship between signal variable and noise variable. Thus, there is no noise-specific effect in the mixed component after denoising by ICA-DP. ICA-DP showed the most powerful denoising performance, which could remove all the noise-specific effects and enhance the signal effects. Fig. 5 shows the signal-and noise-related components extracted by ICA, before and after denoising, when the noise variable is non-linearly related with component #1 and all the components were spatially independent. ICA-SP and ICA-DP could remove the noise effects effectively, while GLM and ComBat could not remove the noise-related component completely

**Fig. 5.**
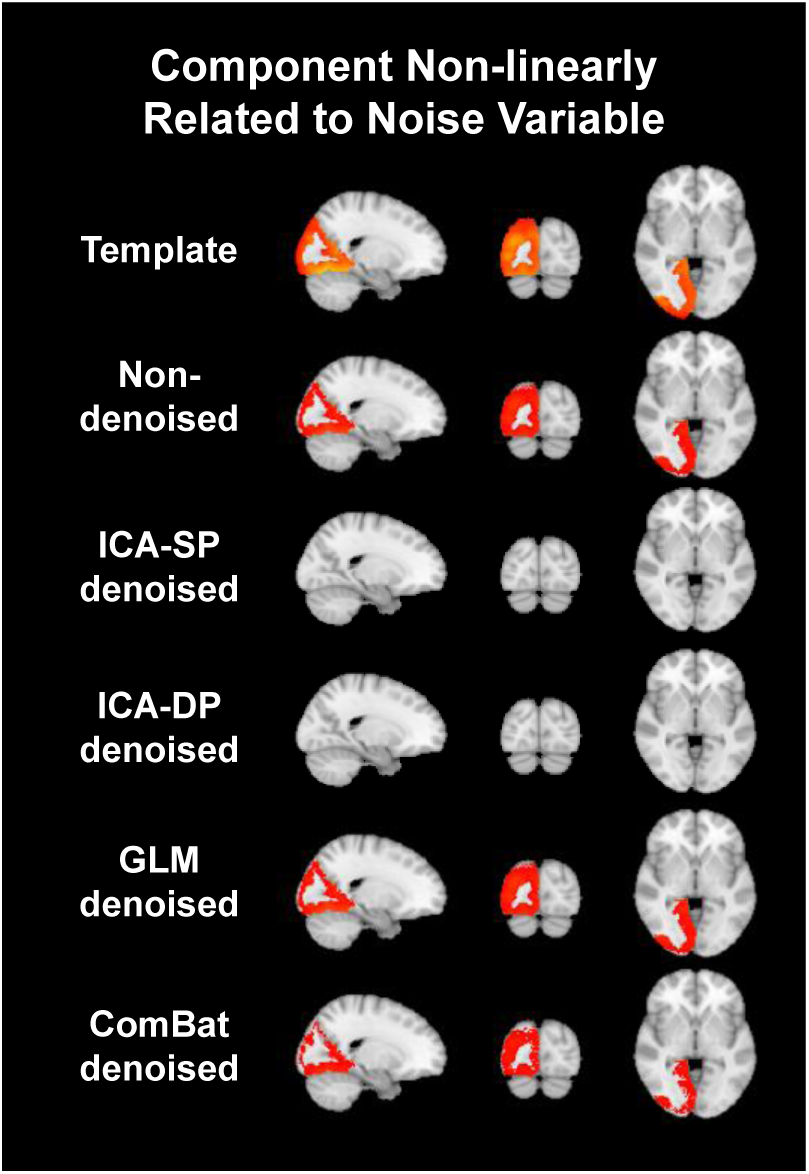
The denoising performance of all the methods when the noise variable is non-linear related with the simulated component #1(ICA was applied to the denoised data to identify the noise-related component). The non-linear correlation coefficients between noise variable and the component #1 extracted from the data denoised by GLM and ComBat are 0.8000 and 0.2099, separately. ICA-based denoising methods show better denoising performance than GLM and ComBat when the site variables have non-linear effects on the data.

when noise variable was non-linear correlated with MRI data. Although ComBat denoising method had not removed the spatial maps of noise-related component completely, it reduced the non-linear correlation coefficient between noise variable and the loading of component #1 from 0.8 to 0.2099 (*p* = 0.0361). The results also confirmed Cetin-Karayumak’s conclusion that ComBat was not good enough to capture and remove the non-linear site noise (Cetin-Karayumak et al. 2020). Besides, GLM did not remove any noise-related information as the non-linear correlation coefficient between the loading of component #1 and noise variable had not changed.

### 3.2 Real Datasets Denoising Results

After denoising, we performed a set of analyses to show the elimination of site effects and the preservation of biological variability, i.e., HC/ASD and age for ABIDE II dataset, and subject heterogeneity for the traveling subject dataset.

Fig. 6 shows the tSNE-2D projection of ADIDE II and traveling subject datasets before and after denoising. The t-SNE was utilized to project the data into two dimensions by using the two dominant features of the non-denoised data and denoised data, to visualize the distribution of site effects and indicate whether it could be eliminated after denoising. For ABIDE II dataset, the data points of the non-denoised data showed site-clustered distribution as most of the centers had their own specific cluster area, except for some intersections among centers UCLA, OHSU, and ETH. And the site-clustered distribution disappeared after being denoised by any of the denoising method. For the traveling subject dataset, the projected data points of the non-denoised data from the same subject tend to be clustered into one cluster, i.e., the first two projected features were dominated by the subject heterogeneity rather than site effects. Though significant difference was not found before and after denoising, the subject heterogeneity was well preserved after denoising.

**Fig. 6.**
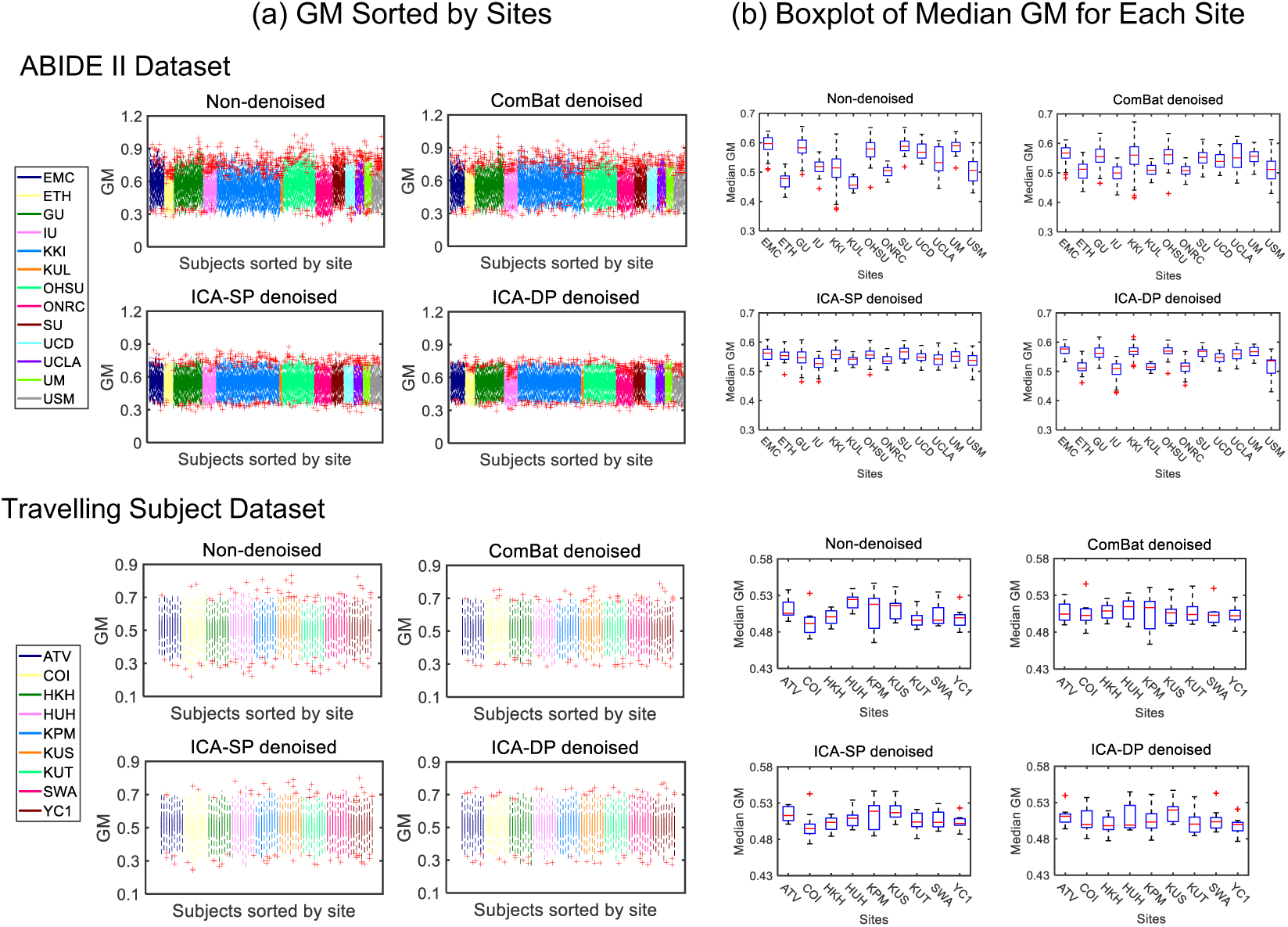
Dimension reduction visualization by t-SNE before and after denoising for ABIDE II and travelling subject datasets. The site-cluster distribution of ABIDE II dataset before denoising indicated the site effects, and it decreased when the data points randomly distributed after denoising. For travelling subject dataset, the subject-cluster distribution indicated the dominant of subject heterogeneity, as the subject from this dataset are the same ones scanned at different center (the data points were labelled by subject numbers). There was no significant difference before and after denoising, and subject heterogeneity was well preserved after denoising.

In Fig. 7(a), diagnostic plots were presented for all the subjects from the two datasets, and the different colors represent different sites. For each subject, the GM measurements were summarized into a boxplot. The different range of GM values among sites was reduced after denoising, and the ICA-based denoising showed efficient reduction. Fig. 7(b) shows the median GM values distribution of the subjects from different sites. After denoising, the site effects decreased noticeably for all the denoising methods, especially for the ICA-based methods denoised data for both ABIDE II and traveling subject datasets.

**Fig. 7.**
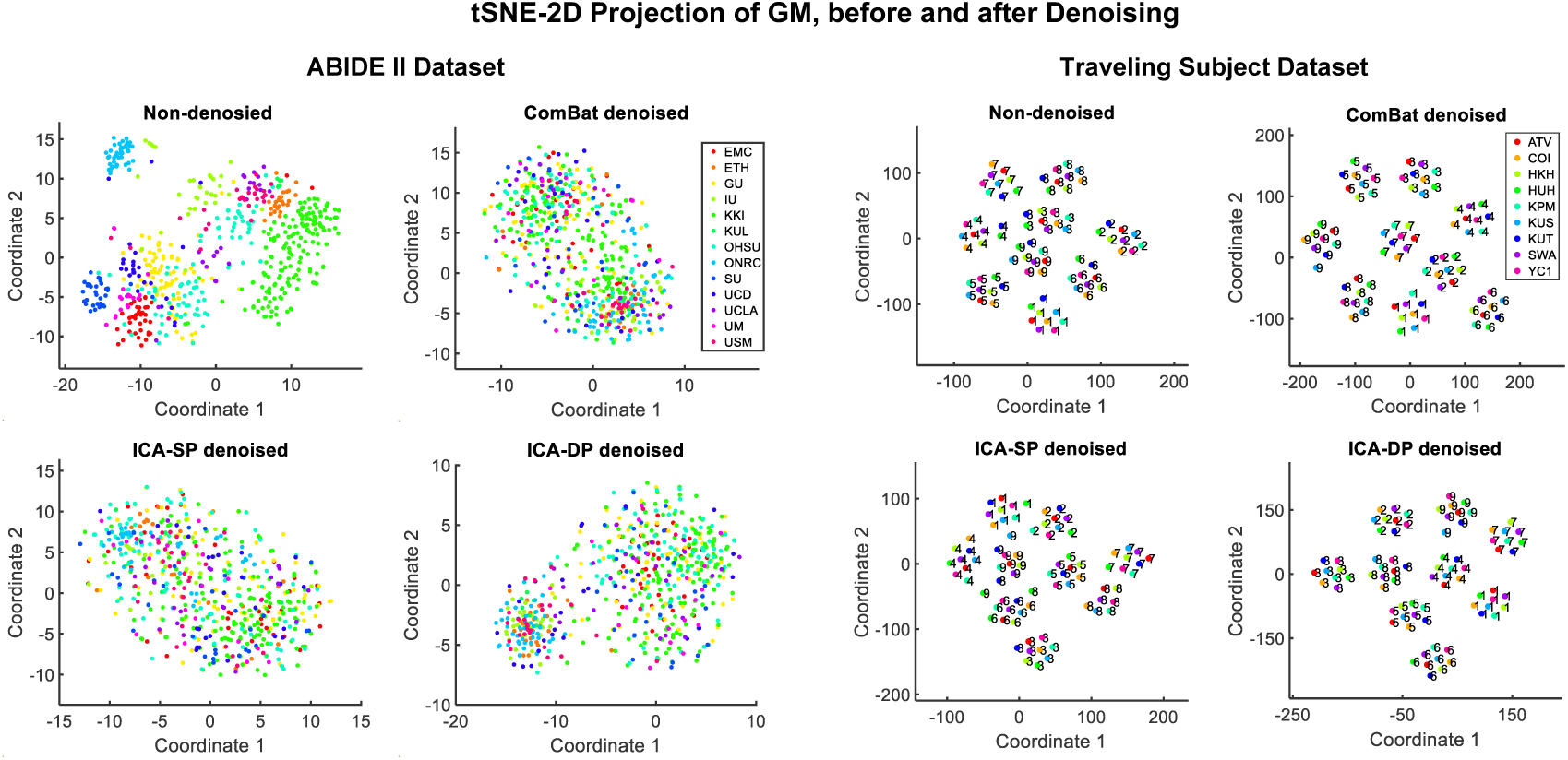
(a) Site-sorted boxplots of GM. Each boxplot represents the GM values distribution of 100 regions of interest (ROI) for every subject. The ranges indicated differences among sites and among subjects. (b) Site-sorted boxplots of median GM. Each boxplot represents the distribution of median GM values for all subjects from the same site. The fluctuates indicated the inter-site difference.

The boxplots presented in Fig.8, for non-denoised data and denoised data, summarized the distribution of the median GM for each subject, which revealed heterogeneity among different subjects. The subject heterogeneity was not destroyed by all the denoising methods. Besides, the intra-subject difference (represented by the height of each box), as a representation of site effects, had been most significantly reduced after denoising by ComBat and ICA-DP.

**Fig. 8.**
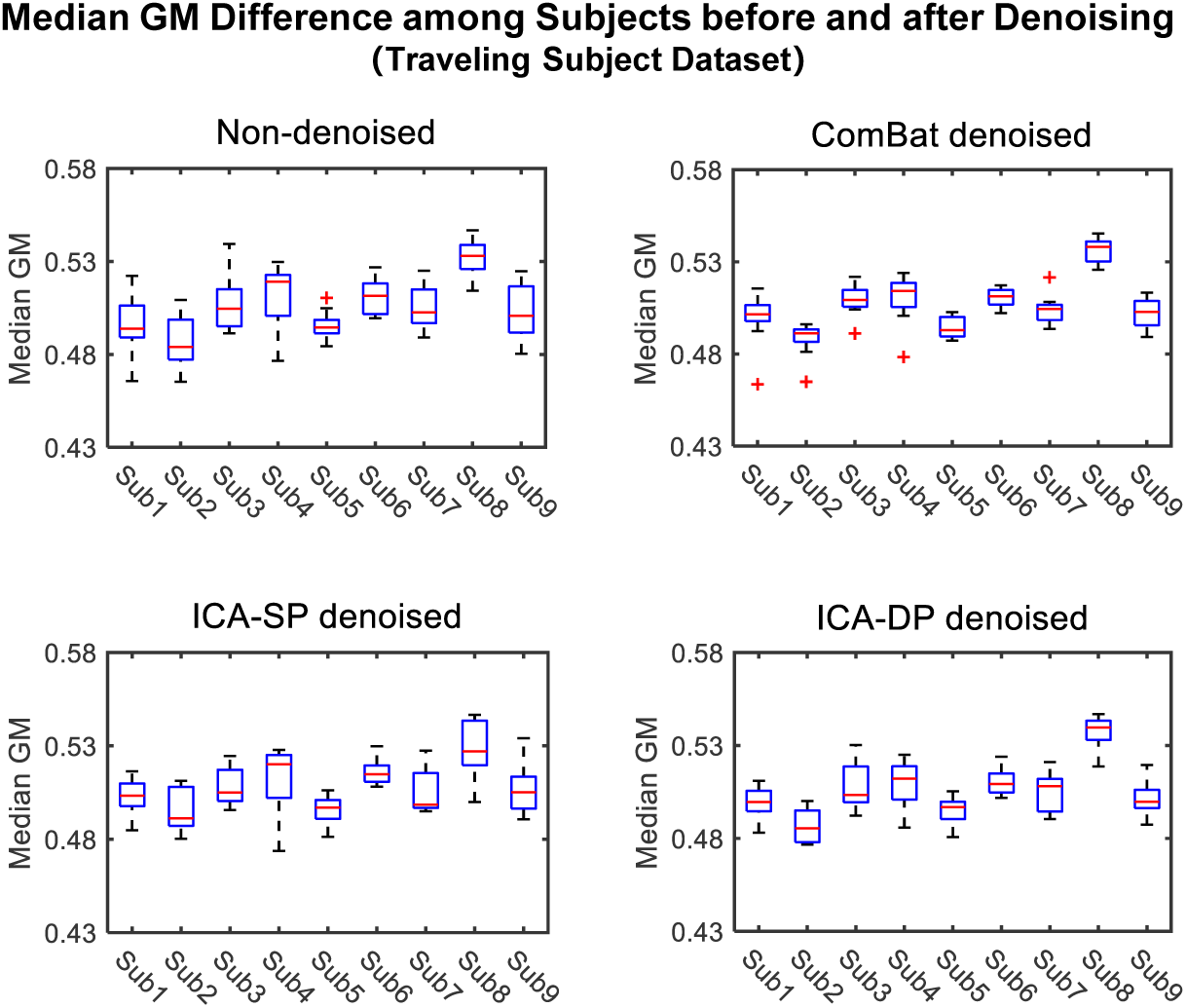
Subject-sorted boxplots of median GM. The fluctuates indicated the inter-subject difference and the height of each box indicated the intra-subject (inter-site) difference.

Fig. 9 shows the group-level analysis for site effects from the two datasets. The non-denoised GM data was globally affected by the site effects for both datasets. Though the site effects had been alleviated by ICA-SP method, it could not remove the effects sufficiently. After denoising by ICA-DP and ComBat, there were no significant regions associated with site variable for both datasets (FWE-corrected *p* < 0.05).

**Fig. 9.**
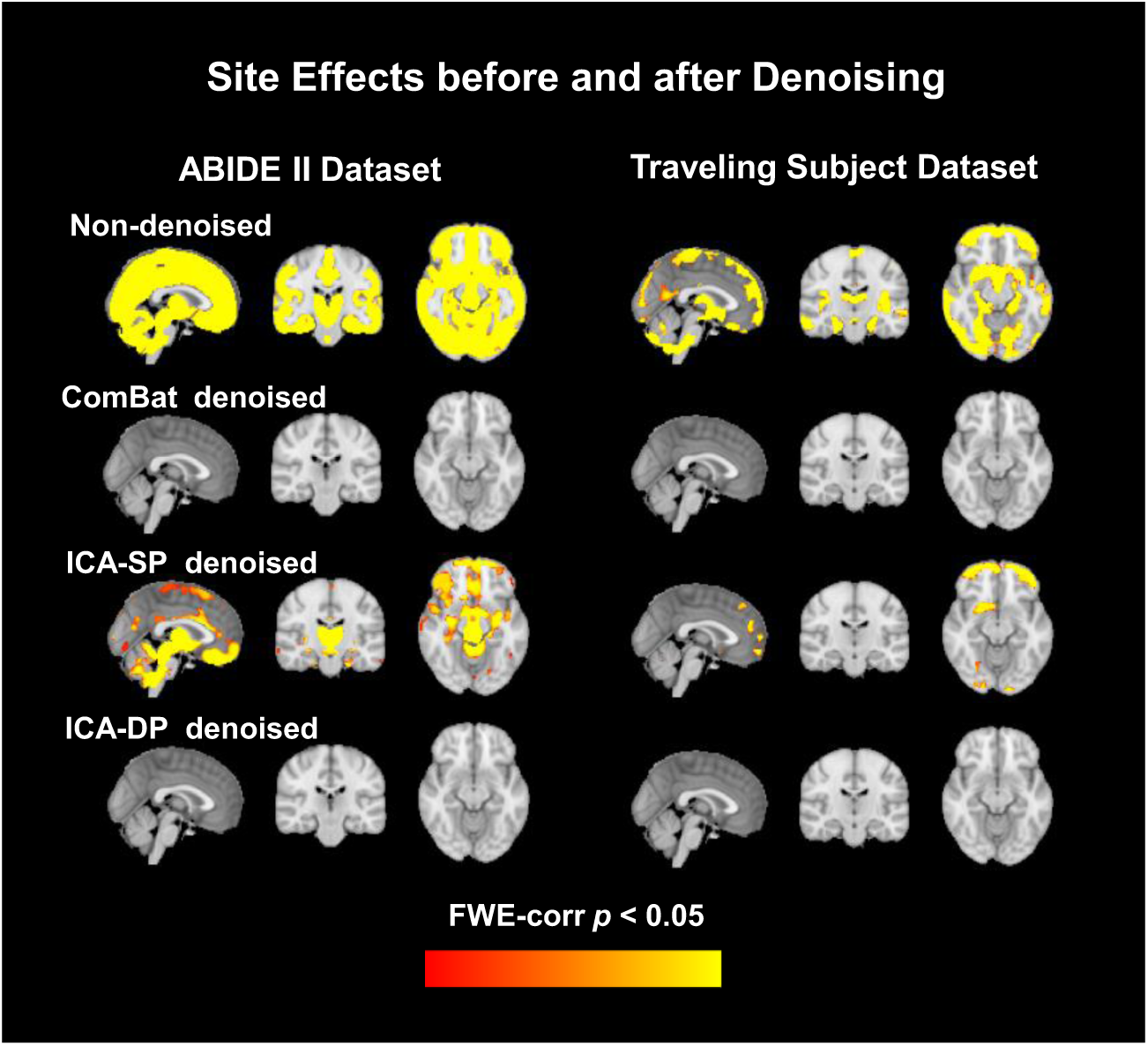
Group-level analysis for site effects before and after denoising. Site effects were removed completely by Combat and ICA-DP. Though ICA-SP reduced the site effects, some significant regions still could be found.

As a biological variable of interest, age effects of the ABIDE II dataset, before and after denoising, were shown in Fig. 10. The correlation between age and median GM before and after denoising with ABIDE dataset were shown in Fig.10(a). The median GM were sorted by age and the data from different scanning centers were in different colors. Pearson correlation coefficients between age and median GM from non-denoised data and denoised data were calculated, which were -0.4746 (Non-denoised), -0.5689 (ComBat denoised), -0.3617 (ICA-SP denoised), -0.8493 (ICA-DP denoised), respectively. The correlation coefficients indicated that the negative correlation between GM and age were strengthened by ICA-DP and ComBat, especially for ICA-DP. Fig.10(b) shows the group-level analyses for age. Site effects confound us to find the true age effects. The negative age effects were not found in the non-denoised data because of the existence of site effects, removal of the effects by all the denoising methods, especially for ICA-DP, could reveal the negative age effects that are not detected from the non-denoised data.

**Fig. 10.**
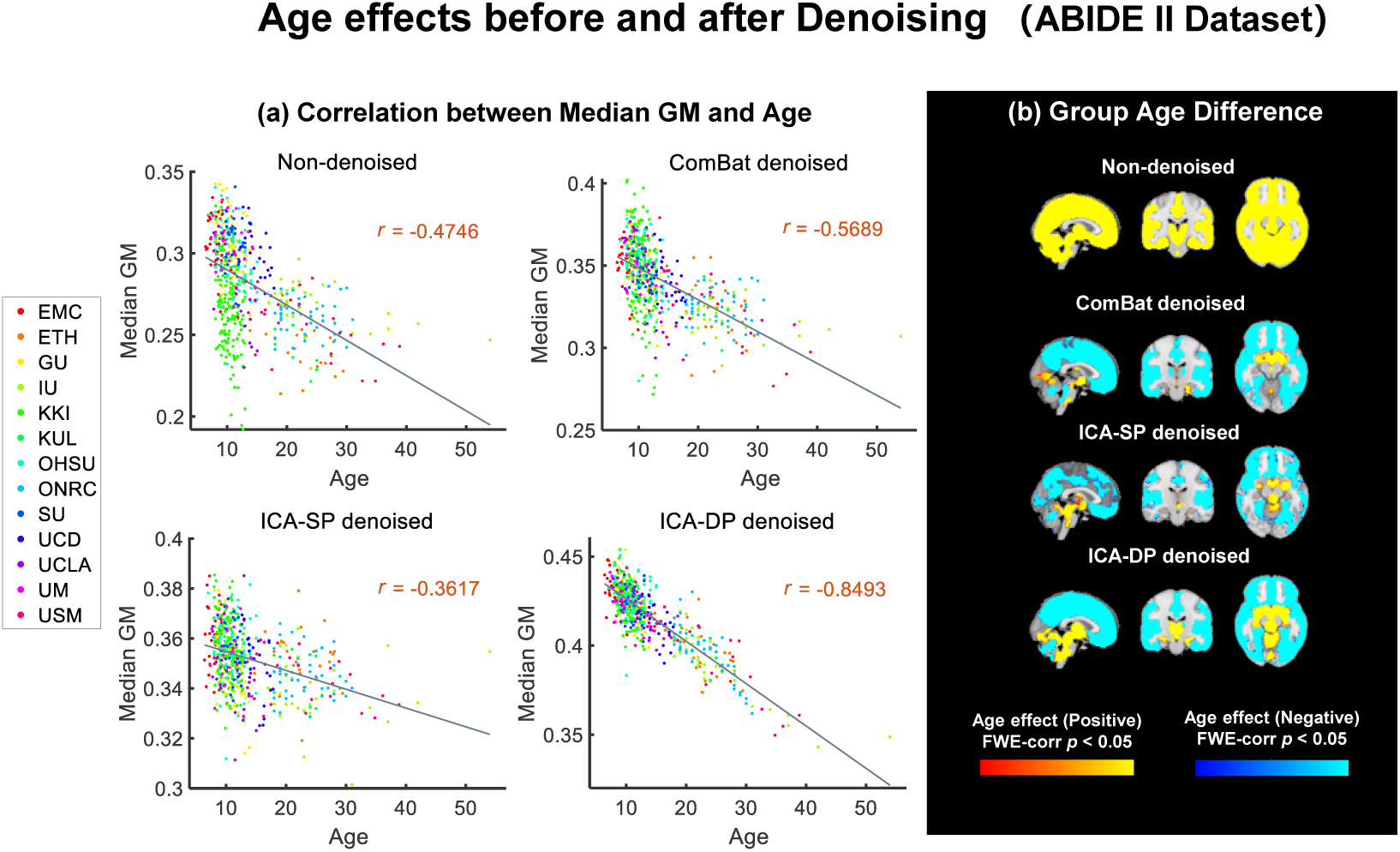
(a) Relationship between age and median GM before and after site-effects denoising. The Pearson correlation coefficient were -0.4746 (Non-denoised), -0.5689 (ComBat denoised), -0.3617 (ICA-SP denoised) and -0.8493 (ICA-DP denoised). (b) Group-level analysis of GM maps for age effects before and after data denoising. The negative age effects were enhanced after denoising, and it could not be found in the non-denoised data when testing age group difference.

Fig. 11 shows the group difference (ASD/HC) before and after denoising. ICA-DP increased the group effects by detecting more significantly different regions related to ASD and HC, while ComBat and ICA-SP decreased the group effects as no significant regions were tested from the data denoised by them. Compared to the non-thresholded group difference maps from non-denoised data (first row), the regions associated with group difference (ASD/HC) from ICA-DP-based denoised data could also be found in the non-denoised data. In other words, ICA-DP only strengthened the signal that should be there rather than reintroducing artifacts.

**Fig. 11.**
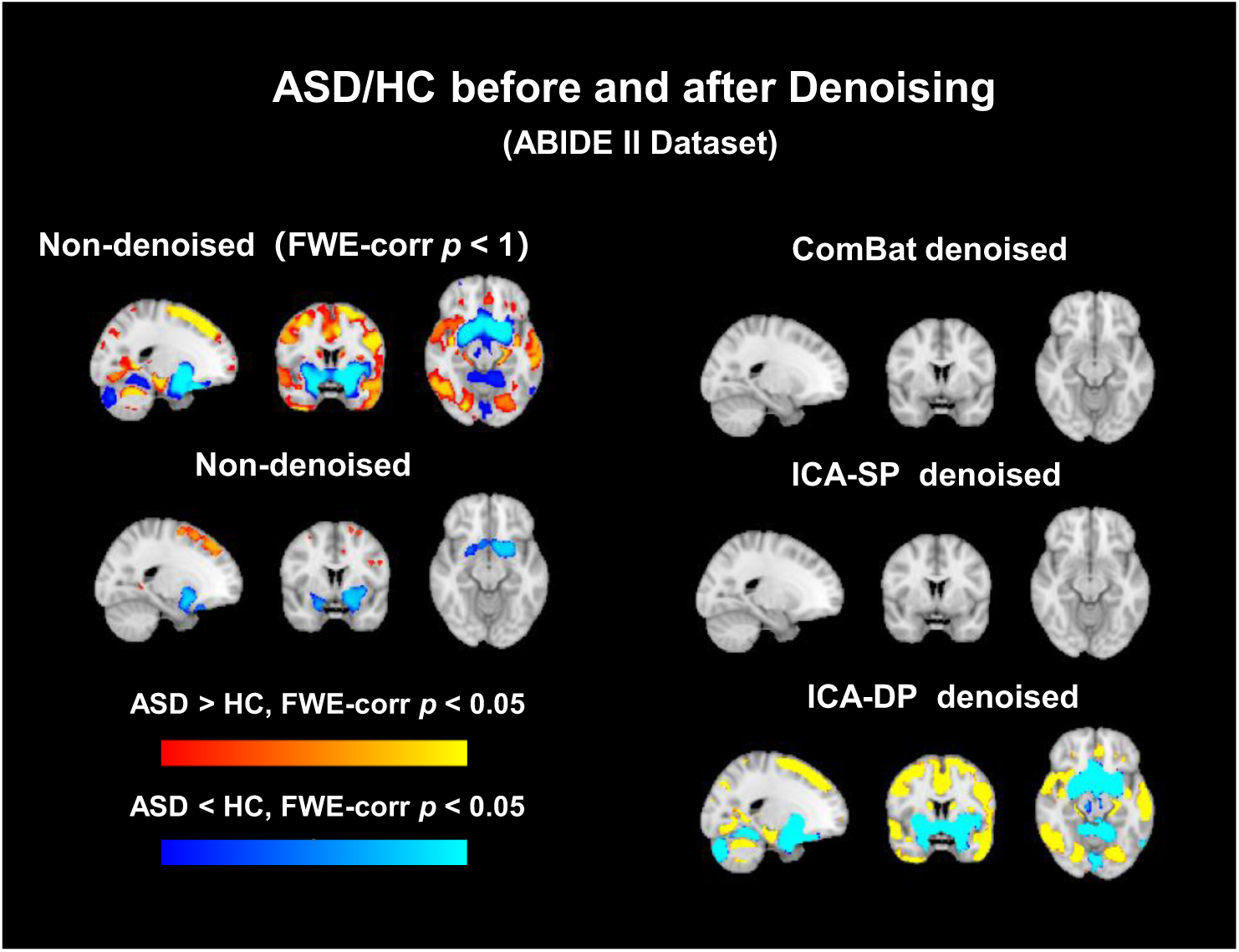
Group-level analysis of GM maps for group difference (ASD/HC) before and after data denoising. No significant regions were found from the data denoised by ComBat and ICA-SP, while ICA-DP could increase the significance of the regions related to ASD/HC. FWE-corr *p* < 1 was shown for non-denoised data to indicate that the regions tested from ICA-DP denoised data were not reintroduced artifacts.

Fig. 12 shows the denoising performance of the two ICA-based denoising methods under different choices of component numbers when running ICA algorithms. The ICA-SP could not remove the site effect completely, though it decreased more site effects as the number of components increasing, and the information related to signal variable (ASD/HC) could not be detected from the data denoised by it under any component number choosing, indicating that this kind of soft denoising based on ICA could neither remove the site effects completely nor reveal the information related to covariates of interest. In contrast, after denoising by ICA-DP, the information that related to site effects could not be tested and there were some regions that significantly correlated to ASD/HC could be revealed from the denoised data, indicating good performance for eliminating site effects and the ability to unveil the signal related information concurrently and showing no affection of which number of components were chosen.

**Fig. 12.**
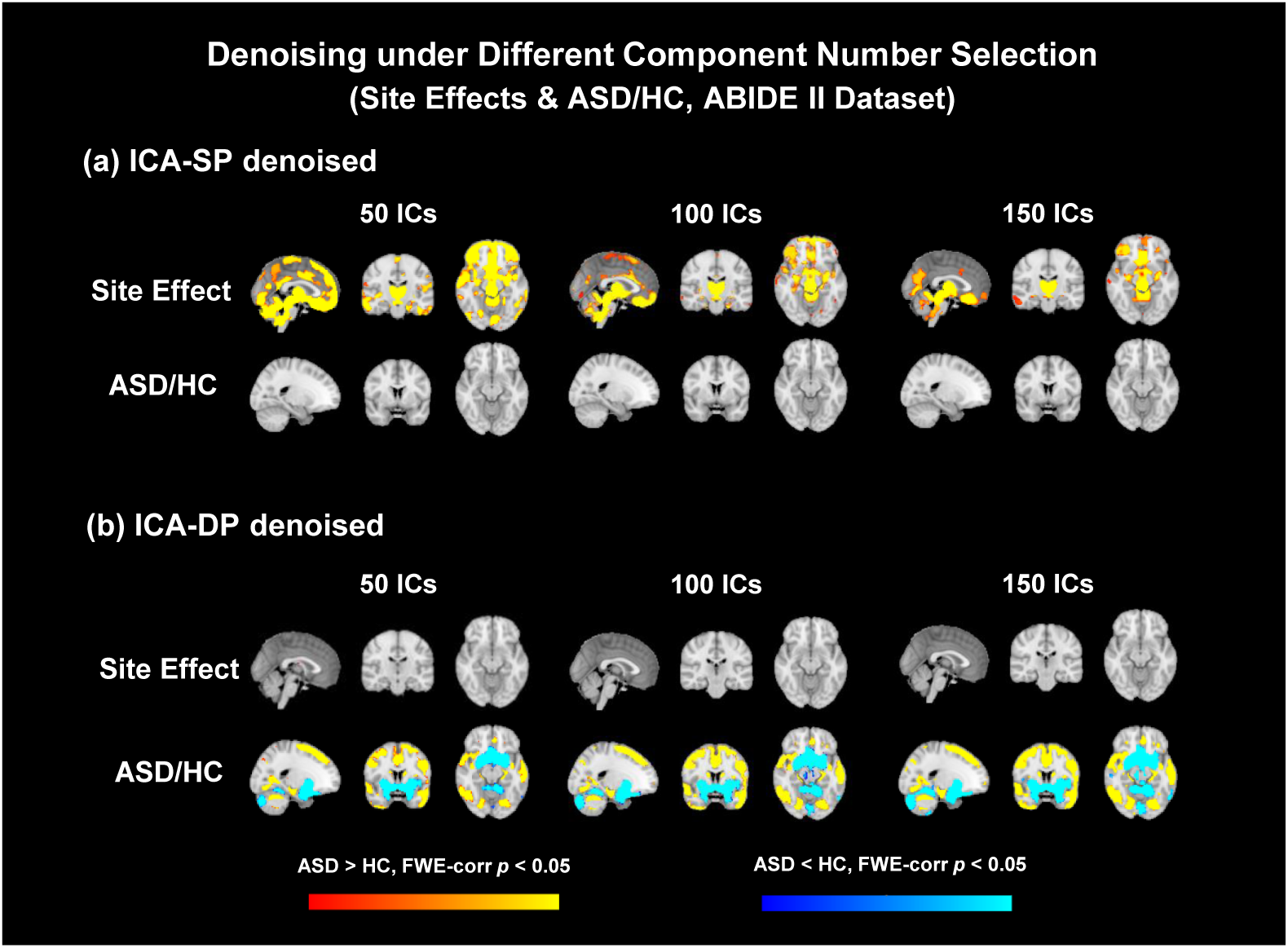
Denoising performance of the two ICA-based denoising methods under different component numbers (50, 100 and 150). After denoising by ICA-DP, site effects were removed completely and ASD/HC group differences were significantly enhanced without the affection of component number.

## 4 Discussion

In this paper, we proposed a dual-projection ICA-based denoising method that can remove the site effects more effectively and completely while enhancing the signal effects. This method shows the superior performance when the site effects are also related with signal variables.

ICA-DP was developed to clean the site effects more completely and effectively when the site effects and signal effects are correlated. Based on the simulation results (Figs. 3, 4, 5), it was found that ICA-DP can eliminate the site effects effectively while enhancing the signal-related information, whether the site effects are linear or non-linear related with MRI data and whether the site and signal variables are correlated. The benefits of ICA-DP are contributed by two reasons: first, ICA-DP can extract all the site-related effects with the first projection even though some site effects are mixed with the signal effects that could not be extracted from original ICA method; second, ICA-DP is more stable than original ICA method, as it can extract and remove all the site effects without the limitation of model order selection; third, compared to standard GLM and ComBat model, scanner-related effects are better captured by ICA-DP, which can capture day-to-day variations in scanner performance. Compared with ICA-DP, ICA-SP, GLM and ComBat showed different defects when denoising the site effects. For ICA-SP, it encountered a problem that it failed to eliminate the site effects completely when the site-related components also significantly correlated to signal variables (Figs. 3, 4). For the GLM-based denoising method, it performed well and was comparable to ICA-DP when the signal and noise variable are not significantly correlated (Fig. 3(a)). However, GLM showed aggressive denoising performance that will remove or decrease the signal-related information when the signal and noise variables are significantly correlated. The signal distortion became worse when the correlation coefficients between the signal and noise variables increased (Fig. 4). Though ComBat-based denoising method showed better performance than GLM and original ICA-based denoising performance which can effectively remove site effects while keeping or strengthening some signal effects when signal and site variables are not correlated. However, ComBat could not completely remove the site effects when signal and site variables are correlated when the mixed components are more correlated with signal (Fig. 4). Meanwhile, we found that ComBat showed aggressive denoising that also removed signal effects when the mixed components are more related with site effects. GLM and ComBat-based method also could not remove the non-linear site effects effectively (Fig. 4).

Based on the results of ABIDE II dataset and the traveling-subject dataset, ICA-DP also shows superior performance on denoising site effects than ICA-SP and ComBat. The significant regions of GM that related to site effects, which cannot be completely removed by ICA-SP, were not detectable after denoising with ICA-DP and ComBat (Fig. 9). The inter-site variation of the traveling data and the intra-subject variation among nine sites for the traveling subject dataset were most significantly reduced after denoising with ICA-DP (Fig. 7). Moreover, subject heterogeneity of the traveling subject dataset was also well preserved after denoising with ICA-DP (Figs. 6, 8).

In addition, ICA-DP also shows superior performance in enhancing biological variability (i.e., age effects and group difference between ASD and HC) compared with ICA-SP and ComBat based on the results of ABIDE II dataset. Site effects hinder us from finding the true age and group effects. The age effects detected with GM of the non-denoised ABIDE II is opposite with the recognized results (Gennatas et al., 2017; Groves et al., 2012). After denoising with ICA-SP, ICA-DP and ComBat, the true age effects on GM were discovered. Among all the three methods, ICA-DP finds the most significant regions related to age and the negative correlation between age and GM was most strongly enhanced by ICA-DP. The relationship of median GM and age was enhanced from -0.4746 (non-denoised) to -0.8493 after denoising with ICA-DP (Fig. 10). In addition, compared to ICA-SP and ComBat, only ICA-DP enhanced the group effects (ASD vs. HC) by detecting more significantly group different regions which cannot be effectively detected by the non-denoised data (Fig. 11). The notable enhancements of the biological variabilities (i.e., age effects and ASD vs. HC) may attribute to the larger proportion of site-related components that we selected for ICA-DP denoising, which could increase the weights of signal we interested and make the signal-related information easier to be detected. On the other hand, this may lead to the other variables which we are not interested in not be well preserved. To protect other variables that we may be interested, we just need to add these variables to **V**_*S*_ in the first projection of ICA-DP denoising method (Eq. (1)). Thus, the ICA-DP is the most effective method for denoising site effects and preserving biological variability among the methods discussed above. Moreover, unlike ICA-SP, the performance of ICA-DP in site effects denoising and signal enhancement were not affected by the number of components chosen for ICA decomposition. It could clean the site effects and strengthen the signal under any selected component number (Fig. 12).

Finally, as a limitation of the proposed denoising method, when the noise variable is strongly related to signal variable (Fig. 4), ICA-DP could not denoise the intersection effects that related with both site and signal variables (neither do other methods except GLM, the most aggressive one that destroying signal-related information severely), thus the denoised data are still correlated to site effects because of the inherent correlation between noise and signal variables (Nevertheless, the correlation values in our simulation is really high and hardly appear in real data study).

Overall, the dual-projection denoising method is more effective and powerful in removing noise effects while preserving signal-related information than other methods mentioned above, and can enhance the sensitivity to detect signal of interest and remove all the noise that are only contributed by site effects. Compared to Combat, it is a data-driven method rather than utilizing the manually designed covariates for regressing. ICA-DP denoising method has great potential for large-scale multi-site studies to produce combined data free from study-site confounds.

## 5 Conclusion

While combing the multi-site MRI data has great convenience that enhances the statistical results and obviates some of the shortcomings of single-site study, the site noise comes naturally, confounding the MRI data analysis and making the results hard to interpret. The traditional methods designed to eliminate the site noise encounter the incomplete or aggressive denoising, i.e., cannot eliminate the noise well or may destroy the signal-related information. To tackle these shortcomings, we proposed a dual-projection data-driven method based on ICA, which can eliminate the noise better and preserve the signal of interest well. And we strongly recommend researchers use the ICA-DP method to denoise the MRI data as it has the ability to extract subject-specific loadings that correspond to the signal or noise variables.

## References

1. Beckmann, C. F., Mackay, C. E., Filippini, N., & Smith, S. M. and others. (2009). Group comparison of resting-state FMRI data using multi-subject ICA and dual regression. NeuroImage, 47(Supp1 1), S148.

2. Bell, A. J., & Sejnowski, T. J. (1995). An information-maximization approach to blind separation and blind deconvolution. Neural Computation, 7(6), 1129–1159.

3. Bell, T. K., Godfrey, K. J., Ware, A. L., Yeates, K. O., & Harris, A. D. (2022). Harmonization of multi-site MRS data with ComBat. NeuroImage, 257, 119330. https://doi.org/10.1016/j.neuroimage.2022.119330

4. Button, K. S., Ioannidis, J. P., Mokrysz, C., Nosek, B. A., Flint, J., Robinson, E. S., & Munafo, M. R. (2013). Power failure: why small sample size undermines the reliability of neuroscience. Nature Reviews Neuroscience, 14(5), 365–376.

5. C Monte-Rubio, G., Segura, B., P Strafella, A., van Eimeren, T., Ibarretxe-Bilbao, N., Diez-Cirarda, M., Eggers, C., Lucas-Jimenez, O., Ojeda, N., & Pena, J. and others. (2022). Parameters from site classification to harmonize MRI clinical studies: Application to a multi-site Parkinson’s disease dataset. Human Brain Mapping.

6. Casey, B. J., Cohen, J. D., O’Craven, K., Davidson, R. J., Irwin, W., Nelson, C. A., Noll, D. C., Hu, X., Lowe, M. J., Rosen, B. R., Truwitt, C. L., & Turski, P. A. (1998). Reproducibility of fMRI results across four institutions using a spatial working memory task. NeuroImage, 8(3), 249–261. https://doi.org/10.1006/nimg.1998.0360

7. Chen, J., Liu, J., Calhoun, V. D., Arias-Vasquez, A., Zwiers, M. P., Gupta, C. N., Franke, B., & Turner, J. A. (2014). Exploration of scanning effects in multi-site structural MRI studies. Journal of Neuroscience Methods, 230, 37–50.

8. Di Martino, A., O’connor, D., Chen, B., Alaerts, K., Anderson, J. S., Assaf, M., Balsters, J. H., Baxter, L., Beggiato, A., & Bernaerts, S. and others. (2017). Enhancing studies of the connectome in autism using the autism brain imaging data exchange II. Scientific Data, 4(1), 1–15.

9. Dinsdale, N. K., Jenkinson, M., & Namburete, A. I. L. (2021). Deep learning-based unlearning of dataset bias for MRI harmonisation and confound removal. NeuroImage, 228(December 2020), 117689. https://doi.org/10.1016/j.neuroimage.2020.117689

10. Eickhoff, S., Nichols, T. E., Van Horn, J. D., & Turner, J. A. (2016). Sharing the wealth: Neuroimaging data repositories. NeuroImage, 124, 1065–1068. https://doi.org/10.1016/j.neuroimage.2015.10.079

11. Fennema-Notestine, C., Gamst, A. C., Quinn, B. T., Pacheco, J., Jernigan, T. L., Thal, L., Buckner, R., Killiany, R., Blacker, D., Dale, A. M., Fischl, B., Dickerson, B., & Gollub, R. L. (2007). Feasibility of multi-site clinical structural neuroimaging studies of aging using legacy data. Neuroinformatics, 5(4), 235–245. https://doi.org/10.1007/s12021-007-9003-9

12. Filippini, N., MacIntosh, B. J., Hough, M. G., Goodwin, G. M., Frisoni, G. B., Smith, S. M., Matthews, P. M., Beckmann, C. F., & Mackay, C. E. (2009). Distinct patterns of brain activity in young carriers of the APOE-$\varepsilon$4 allele. Proceedings of the National Academy of Sciences, 106(17), 7209–7214.

13. Focke, N. K., Helms, G., Kaspar, S., Diederich, C., Tóth, V., Dechent, P., Mohr, A., & Paulus, W. (2011). Multi-site voxel-based morphometry-Not quite there yet. NeuroImage, 56(3), 1164–1170. https://doi.org/10.1016/j.neuroimage.2011.02.029

14. Fortin, J. P., Parker, D., Tunç, B., Watanabe, T., Elliott, M. A., Ruparel, K., Roalf, D. R., Satterthwaite, T. D., Gur, R. C., Gur, R. E., Schultz, R. T., Verma, R., & Shinohara, R. T. (2017). Harmonization of multi-site diffusion tensor imaging data. NeuroImage, 161(July), 149–170. https://doi.org/10.1016/j.neuroimage.2017.08.047

15. Friedman, L., Stern, H., Brown, G. G., Mathalon, D. H., Turner, J., Glover, G. H., Gollub, R. L., Lauriello, J., Lim, K. O., Cannon, T., Greve, D. N., Bockholt, H. J., Belger, A., Mueller, B., Doty, M. J., He, J., Wells, W., Smyth, P., Pieper, S., … Potkin, S. G. (2008). Test-retest and between-site reliability in a multicenter fMRI study. Human Brain Mapping, 29(8), 958–972. https://doi.org/10.1002/hbm.20440

16. Gennatas, E. D., Avants, B. B., Wolf, D. H., Satterthwaite, T. D., Ruparel, K., Ciric, R., Hakonarson, H., Gur, R. E., & Gur, R. C. (2017). Age-related effects and sex differences in gray matter density, volume, mass, and cortical thickness from childhood to young adulthood. Journal of Neuroscience, 37(20), 5065–5073.

17. Glover, G. H., Mueller, B. A., Turner, J. A., Van Erp, T. G. M., Liu, T. T., Greve, D. N., Voyvodic, J. T., Rasmussen, J., Brown, G. G., Keator, D. B., Calhoun, V. D., Lee, H. J., Ford, J. M., Mathalon, D. H., Diaz, M., O’Leary, D. S., Gadde, S., Preda, A., Lim, K. O., … Potkin, S. G. (2012). Function biomedical informatics research network recommendations for prospective multicenter functional MRI studies. Journal of Magnetic Resonance Imaging, 36(1), 39–54. https://doi.org/10.1002/jmri.23572

18. Groves, A. R., Beckmann, C. F., Smith, S. M., & Woolrich, M. W. (2011). Linked independent component analysis for multimodal data fusion. Neuroimage, 54(3), 2198–2217.

19. Groves, A. R., Smith, S. M., Fjell, A. M., Tamnes, C. K., Walhovd, K. B., Douaud, G., Woolrich, M. W., & Westlye, L. T. (2012). Benefits of multi-modal fusion analysis on a large-scale dataset: life-span patterns of inter-subject variability in cortical morphometry and white matter microstructure. Neuroimage, 63(1), 365–380.

20. Johnson, W. E., Li, C., & Rabinovic, A. (2007). Adjusting batch effects in microarray expression data using empirical Bayes methods. Biostatistics, 8(1), 118–127. https://doi.org/10.1093/biostatistics/kxj037

21. Jovicich, J., Czanner, S., Han, X., Salat, D., Kouwe, A. Van Der Quinn, B., Pacheco, J., Albert, M., Killiany, R., Blacker, D., Rosas, D., Makris, N., Gollub, R., & Dale, A. (2009). MRI-derived measurements of human subcortical, ventricular andintracranial brain volumes: Reliability effects of scan sessions,acquisition sequences, data analyses, scanner upgrade, scannervendors and field strengths. NeuroImage, 46(1), 177–192. https://doi.org/10.1016/j.neuroimage.2009.02.010.MRI-derived

22. Li, H., Smith, S. M., Gruber, S., Lukas, S. E., Silveri, M. M., Hill, K. P., Killgore, W. D., & Nickerson, L. D. (2020). Denoising scanner effects from multimodal MRI data using linked independent component analysis. Neuroimage, 208, 116388.

23. Maikusa, N., Zhu, Y., Uematsu, A., Yamashita, A., Saotome, K., Okada, N., Kasai, K., Okanoya, K., Yamashita, O., & Tanaka, S. C. and others. (2021). Comparison of traveling-subject and ComBat harmonization methods for assessing structural brain characteristics. Human Brain Mapping, 42(16), 5278–5287.

24. Nickerson, L. D., Smith, S. M., {\“O}ng{\”u}r, D., & Beckmann, C. F. (2017). Using dual regression to investigate network shape and amplitude in functional connectivity analyses. Frontiers in Neuroscience, 11, 115.

25. Orlhac, F., Boughdad, S., Philippe, C., Stalla-Bourdillon, H., Nioche, C., Champion, L., Soussan, M., Frouin, F., Frouin, V., & Buvat, I. (2018). A postreconstruction harmonization method for multicenter radiomic studies in PET. Journal of Nuclear Medicine, 59(8), 1321–1328. https://doi.org/10.2967/jnumed.117.199935

26. Pohl, K. M., Sullivan, E. V, Rohlfing, T., Chu, W., Kwon, D., Nichols, B. N., Zhang, Y., Brown, S. A., Tapert, S. F., Cummins, K., Thompson, W. K., Brumback, T., Colrain, I. M., Baker, F. C., Prouty, D., De Bellis, M. D., Voyvodic, J. T., Clark, D. B., Schirda, C., … Pfefferbaum, A. (2016). Harmonizing DTI measurements across scanners to examine the development of white matter microstructure in 803 adolescents of the NCANDA study. NeuroImage, 130, 194–213. https://doi.org/10.1016/j.neuroimage.2016.01.061

27. Smith, S. M., & Nichols, T. E. (2009). Threshold-free cluster enhancement: addressing problems of smoothing, threshold dependence and localisation in cluster inference. Neuroimage, 44(1), 83–98.

28. Takao, H., Hayashi, N., & Ohtomo, K. (2011). Effect of scanner in longitudinal studies of brain volume changes. Journal of Magnetic Resonance Imaging, 34(2), 438–444. https://doi.org/10.1002/jmri.22636

29. Tanaka, S. C., Yamashita, A., Yahata, N., Itahashi, T., Lisi, G., Yamada, T., Ichikawa, N., Takamura, M., Yoshihara, Y., & Kunimatsu, A. and others. (2021). A multi-site, multi-disorder resting-state magnetic resonance image database. Scientific Data, 8(1), 1–15.

30. Tian, D., Zeng, Z., Sun, X., Tong, Q., Li, H., He, H., Gao, J. H., He, Y., & Xia, M. (2022). A deep learning-based multisite neuroimage harmonization framework established with a traveling-subject dataset. NeuroImage, 257(March), 119297. https://doi.org/10.1016/j.neuroimage.2022.119297

31. Van Horn, J. D., & Toga, A. W. (2009). Multisite neuroimaging trials. Current Opinion in Neurology, 22(4), 370–378. https://doi.org/10.1097/WCO.0b013e32832d92de

32. Venkatraman, V. K., Gonzalez, C. E., Landman, B., Goh, J., Reiter, D. A., An, Y., & Resnick, S. M. (2015). Region of interest correction factors improve reliability of diffusion imaging measures within and across scanners and field strengths. Neuroimage, 119, 406–416.

33. Vollmar, C., O’Muircheartaigh, J., Barker, G. J., Symms, M. R., Thompson, P., Kumari, V., Duncan, J. S., Richardson, M. P., & Koepp, M. J. (2010). Identical, but not the same: Intra-site and inter-site reproducibility of fractional anisotropy measures on two 3.0T scanners. NeuroImage, 51(4), 1384–1394. https://doi.org/10.1016/j.neuroimage.2010.03.046

34. Wegner, C., Filippi, M., Korteweg, T., Beckmann, C., Ciccarelli, O., De Stefano, N., Enzinger, C., Fazekas, F., Agosta, F., Gass, A., Hirsch, J., Johansen-Berg, H., Kappos, L., Barkhof, F., Polman, C., Mancini, L., Manfredonia, F., Marino, S., Miller, D. H., … Matthews, P. M. (2008). Relating functional changes during hand movement to clinical parameters in patients with multiple sclerosis in a multi-centre fMRI study. European Journal of Neurology, 15(2), 113–122. https://doi.org/10.1111/j.1468-1331.2007.02027.x

35. Winkler, A. M., Ridgway, G. R., Webster, M. A., Smith, S. M., & Nichols, T. E. (2014). Permutation inference for the general linear model. Neuroimage, 92, 381–397.

36. Yamashita, A., Yahata, N., Itahashi, T., Lisi, G., Yamada, T., Ichikawa, N., Takamura, M., Yoshihara, Y., Kunimatsu, A., Okada, N., Yamagata, H., Matsuo, K., Hashimoto, R., Okada, G., Sakai, Y., Morimoto, J., Narumoto, J., Shimada, Y., Kasai, K., … Imamizu, H. (2019). Harmonization of resting-state functional MRI data across multiple imaging sites via the separation of site differences into sampling bias and measurement bias. In PLoS Biology. https://doi.org/10.1101/440875

37. Yu, M., Linn, K. A., Cook, P. A., Phillips, M. L., McInnis, M., Fava, M., Trivedi, M. H., Weissman, M. M., Shinohara, R. T., & Sheline, Y.I. (2018). Statistical harmonization corrects site effects in functional connectivity measurements from multi-site fMRI data. Human Brain Mapping, 39(11), 4213– 4227. https://doi.org/10.1002/hbm.24241

38. Zivadinov, R., & Cox, J. L. (2008). Is functional MRI feasible for multi-center studies on multiple sclerosis? European Journal of Neurology, 15(2), 109–110. https://doi.org/10.1111/j.1468-1331.2007.02030.x

